# Cell competition acts as a purifying selection to eliminate cells with mitochondrial defects during early mouse development

**DOI:** 10.1101/2020.01.15.900613

**Authors:** Ana Lima, Gabriele Lubatti, Jörg Burgstaller, Di Hu, Alistair Green, Aida Di Gregorio, Tamzin Zawadzki, Barbara Pernaute, Elmir Mahammadov, Salvador Perez Montero, Marian Dore, Juan Miguel Sanchez, Sarah Bowling, Margarida Sancho, Mohammad Karimi, David Carling, Nick Jones, Shankar Srinivas, Antonio Scialdone, Tristan A. Rodriguez

## Abstract

Cell competition is emerging as a quality control mechanism that eliminates unfit cells in a wide range of settings from development to the adult. However, the nature of the cells normally eliminated by cell competition and what triggers their elimination remains poorly understood. In mouse, prior to gastrulation 35% of epiblast cells are eliminated. Here we have performed single cell transcriptional profiling of these cells and find that they show the hallmarks of cell competition and have mitochondrial defects. We demonstrate that mitochondrial defects are common to a range of different loser cell types and that manipulating mitochondrial function is sufficient to trigger competition. Importantly, we show that in the embryo cell competition eliminates cells with mitochondrial DNA mutations and that even non-pathological changes in mitochondrial DNA sequence can induce cell competition. Our results therefore suggest that cell competition is a purifying selection that optimises mitochondrial performance prior to gastrulation.

---

Cell competition is a fitness sensing mechanism that eliminates cells that, although viable, are less fit than their neighbours. The cells that are eliminated are generically termed losers, while the fitter cells that survive are referred to as winners. Cell competition has been shown to act in a broad range of settings, from the developing embryo to the ageing organisms^1–3^. It has been primarily studied in *Drosophila*, where it was first described in the imaginal wing disc ^4^. Since then, it has also been found to be conserved in mammals. In the mouse embryo 35% of embryonic cells are eliminated between E5.5 and E6.5 and there is strong evidence that this elimination is through cell competition^5–7^. These and other studies identified a number of read-outs of cell competition in the mouse embryo, such as relative low c-MYC expression, a loss of mTOR signalling, low TEAD activity, high P53 expression, or elevated levels of ERK phosphorylation ^5–9^. Importantly, there is a significant overlap with the markers of cell competition originally identified in *Drosophila* as well as those found in other cell competition models, such as Madin-Darby Canine Kidney (MDCK) cells – as reviewed^1–3^. In spite of the advance that having these cell competition markers signifies, given that they were primarily identified by using genetic models that rely on over-expression or mutation, we still have little insight into the over-arching features of the cells that are eliminated in the physiological context.

Mitochondria, with their diverse cellular functions ranging from determining the bioenergetic output of the cell to regulating its apoptotic response, are strong candidates for determining competitive cell fitness. During early mouse development mitochondria undergo profound changes in their shape and activity^10^. In the pre-implantation embryo mitochondria are rounded, fragmented and contain sparse cristae, but upon implantation they fuse to form complex networks with mature cristae^11^. The mode of replication of the mitochondrial genome (mtDNA), that encodes for vital components of the bioenergetic machinery, also changes during early mouse development. After fertilization, mtDNA replication ceases and its copy number per cell decreases with every division until post-implantation stages, when mtDNA replication resumes^10^. As the mutation rate of mtDNA is significantly higher than that of nuclear DNA^12, 13^, this increased replication most likely leads to an increased mutation load. In fact, inheritable mtDNA based diseases are reported with a prevalence of 5-15 cases per 100,000 individuals^14, 15^. A number of mechanisms have been proposed to reduce this mutation load, such as the bottleneck effect, purifying selection or biased segregation of mtDNA haplotypes^16–21^. However, how these mechanisms act at the molecular and cellular level is still poorly understood.

To understand the nature of the cells eliminated during early mouse post-implantation development, we have analysed their transcriptional profile by single-cell RNA sequencing and found that these cells share a cell competition signature. Analysis of the pathways mis-regulated identified mitochondrial dysfunction as a common feature. Importantly, our studies uncovered that the cells eliminated have mtDNA mutations. Furthermore, we demonstrate that manipulating mitochondrial activity either by disrupting mitochondrial dynamics or by introducing non-pathological mtDNA changes is sufficient to trigger cell competition. These results therefore pinpoint mitochondrial performance as a key cellular feature that determines the competitive ability of embryonic cells and suggest that cell competition is acting as a purifying selection during early mammalian development.

## Results

### Cells eliminated in the early mouse embryo have a distinct transcriptional profile

We have previously shown that in the early post-implantation mouse embryo about 35% of epiblast cells are eliminated and that these cells are marked by low mTOR signalling^7^. However, we currently do not understand the characteristics of these cells or what triggers their elimination. To answer these questions, we have analysed their transcriptional profile by single cell RNA sequencing (scRNA-seq). To ensure we can capture the eliminated cells, as we have done before^7^, we isolated embryos at E5.5 and cultured them for 16 hours in the presence of a caspase inhibitors (CI) or vehicle (DMSO) (Fig. 1a). Unsupervised clustering of the scRNA-seq data revealed five clusters: two corresponding to extra-embryonic tissues (visceral endoderm and extra-embryonic ectoderm) and three that expressed epiblast marker genes (Fig. 1b-c, Extended Data Fig. 1a-f and Methods). Interestingly, cells from CI- and DMSO-treated embryos are unequally distributed across the three epiblast clusters. In particular, one of these clusters (cluster 4) is only composed of cells from CI-treated embryos (Fig. 1d-e). It is worth noting that all epiblast clusters contained cells in G2/M and S phases of the cell cycle, suggesting they are all cycling (Extended Data Fig. 2a).

**Fig. 1.**
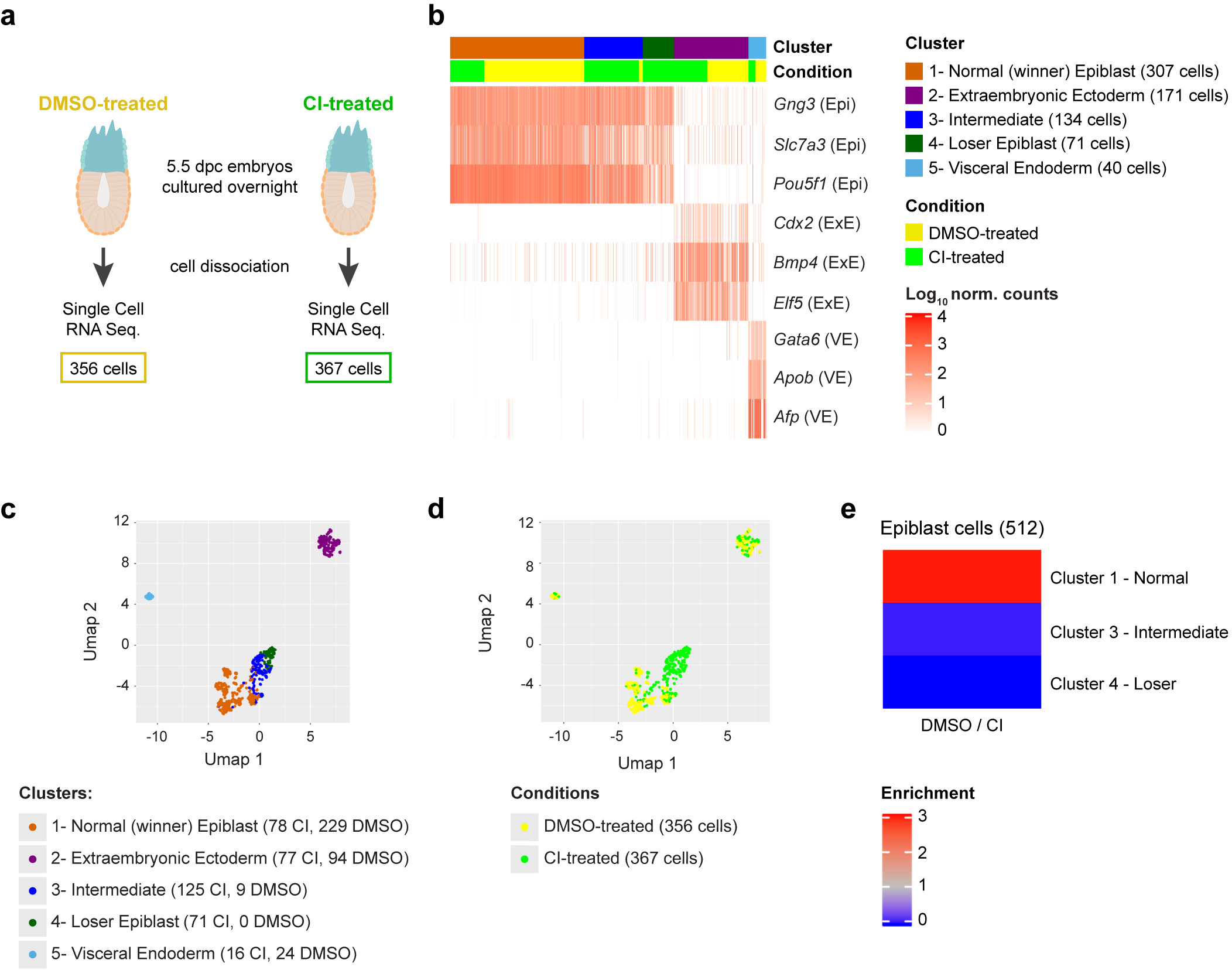
Cells eliminated during early mouse embryogenesis have a distinct transcriptional profile. **a,** Experimental design. The number of cells in the two conditions (DMSO-treated and CI-treated) refers to the number of cells that passed the quality control. **b**, Identification of the clusters according to known gene markers from the different embryonic regions^56^. Three clusters (clusters 1, 3 and 4) show marker genes of the epiblast (Epi), while the remaining clusters correspond to the extra-embryonic lineages visceral endoderm (VE; cluster 5) and extraembryonic ectoderm (ExE; cluster 2). The epiblast clusters are named “Winner”, “Intermediate” and “Loser” on the basis of the relative fraction of cells from CI-treated embryos they include (see panel **e**). **c**,**d**, UMAP visualization of the single-cell RNA-seq data, with cells coloured according to cluster (**c**) or condition (**d**). A region made up exclusively by cells from CI-treated embryos emerges. **e**, Ratio between the fraction of cells from DMSO-treated and CI-treated embryos in the three epiblast clusters. While the “winner” epiblast cluster shows an enrichment of cells from DMSO-treated embryos, the “intermediate” and the “loser” epiblast clusters are strongly enriched for cells from CI-treated embryos.

The three epiblast clusters are highly connected, as highlighted by a connectivity analysis carried out with PAGA^22^ (Extended Data Fig. 2b). Hence, to establish the relationship between these epiblast clusters we computed a diffusion map^23^. For this, we selected only cells captured from CI-treated embryos, to eliminate possible confounding effects due to the caspase inhibitor (Fig. 2a). However, when all epiblast cells are considered, the results remain unchanged (Extended Data Fig. 2c-e). This analysis identified a trajectory between the three epiblast clusters, with those cells unique to CI-treated embryos falling at one extreme end of the trajectory (corresponding to cluster 4; Fig. 2a) and with those cells present in both DMSO and CI-treated embryos at the other (corresponding to cluster 1; Fig. 2a and Extended Data Fig. 2d).

**Fig. 2.**
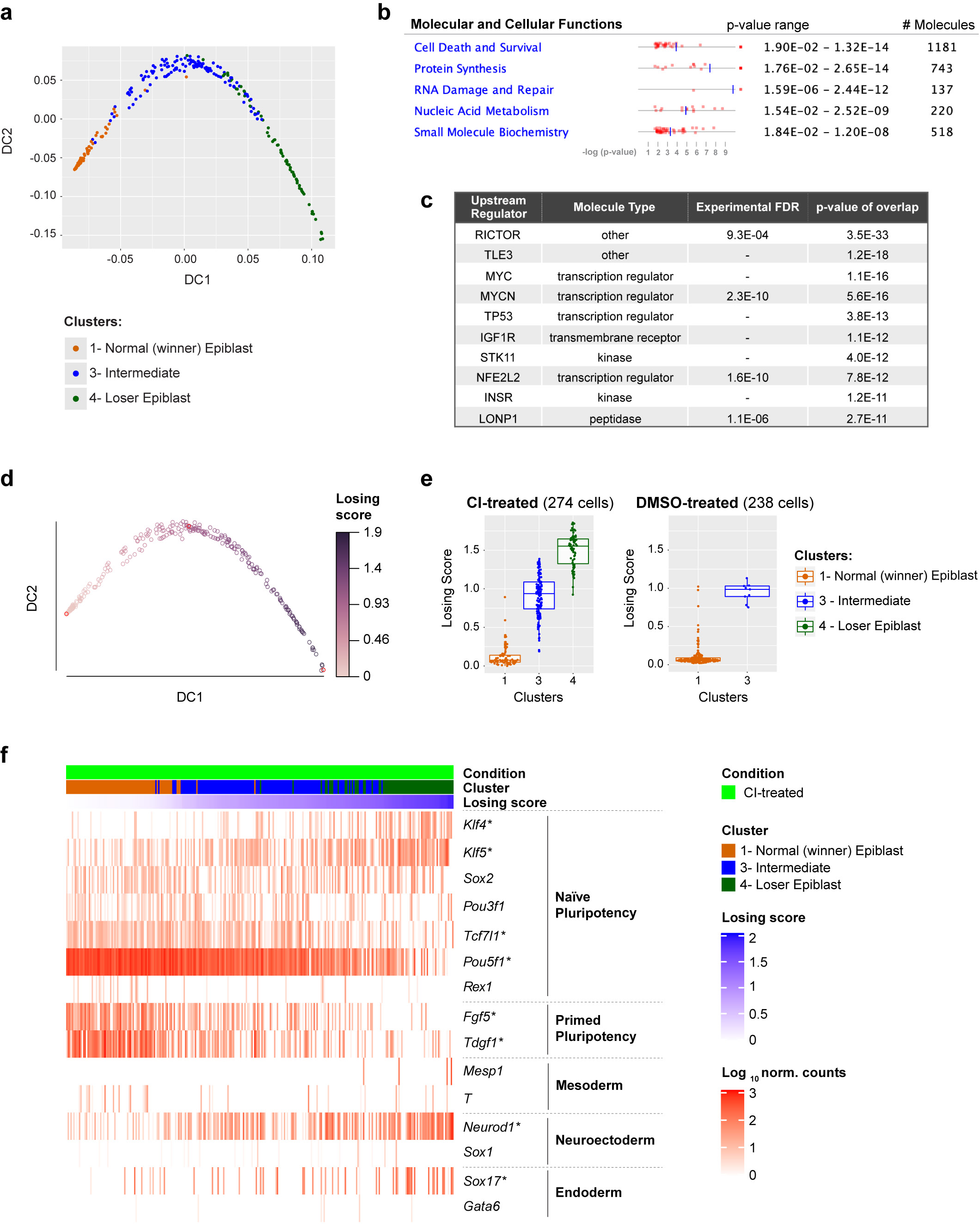
A cell competition transcriptional signature is identified in cells eliminated during mouse embryonic development. **a**, Diffusion map of epiblast cells (only from CI-treated embryos), coloured by cluster. **b**, **c**, IPA run on the list of genes differentially expressed along the diffusion trajectory (see Extended Data Fig. 2a) generated lists of top 5 molecular and cellular functions (**b**) and upstream regulators (**c**) found to be differentially activated in epiblast cells along the diffusion trajectory from winner (cluster 1) to loser status (cluster 4). **d**, Diffusion map of epiblast cells (only from CI-treated embryos) coloured by diffusion pseudotime coordinate (dpt). The winner and the loser clusters are found at the two extremities of the trajectory, hence the dpt can be interpreted as a “losing score”. **e**, Losing score of the cells in the three epiblast clusters in CI-treated (left) or DMSO-treated (right) embryos. The losing score of the cells from DMSO-treated embryos was obtained by projecting them on the diffusion map shown in panel **d** (see Methods). **f,** Expression levels in epiblast cells from CI-treated embryos of genes (in rows) that are markers for naïve pluripotency (*Klf4*, *Klf5, Sox2*, *Pou3f1*, *Tcf7l1* and *Pou5f1 and Rex1*), primed pluripotency (*Fgf5* and *Tdgf1*), mesoderm (*Mesp1* and *T*), neuroectoderm (*Neurod1* and *Sox1*) and endoderm (*Sox17 and Gata6*). Cells (in columns) are sorted by their losing scores. The genes marked with a * are differentially expressed along the trajectory.

To further define the identity of the epiblast cells of CI-treated embryos we analysed the genes differentially expressed along the trajectory (see Methods and Extended Data Fig. 3a) using Ingenuity Pathway Analysis (IPA) to characterize gene signatures^24^. Importantly, we found that these differentially expressed genes fell under molecular and cellular function categories associated with cell death and survival, protein synthesis and nucleic acids (Fig. 2b). Analysis of the factors with enriched targets within the genes differentially expressed along the trajectory revealed RICTOR (an mTOR component), TLE3, MYC, MYCN, P53 and IGFR (that is upstream of mTOR) as the top upstream regulators (Fig. 2c). Breaking down the differentially expressed genes into those down-regulated or up-regulated along the winner-to loser trajectory revealed that the targets of RICTOR, MYC, MYCN and IGFR primarily fell within the down-regulated genes (Supplementary Tables 1 and 2). P53 activated targets were preferentially up-regulated and P53 repressed targets were preferentially down-regulated (Extended Data Fig. 3b-c).

Moreover, genes related to protein synthesis were primarily found to be downregulated.

The observation that the genes differentially expressed along the trajectory fall into cell death categories, as well as being mTOR, MYC and P53 targets strongly suggests that cells at each end of the trajectory are the winners and losers of cell competition^5–7^. For this reason, we hereafter refer to those epiblast cells unique to CI-treated embryos as “loser” epiblast cells and to those at the opposite end of the trajectory as the “winner” epiblast cells. Those cells lying between these two populations on the trajectory are considered “intermediate”. Using this knowledge we can define a diffusion pseudotime (dpt) coordinate^25^ originating in the “winner” cluster that tracks the position of cells along the trajectory and that can be interpreted as a “losing score”, i.e., it quantifies how strong the signature of the “losing” state is in the transcriptome of a cell (see Fig. 2d-e).

In accordance with previous studies^6, 8, 9^, we also found evidence for miss-patterning in the eliminated epiblast cells, as a proportion of these cells co-expressed naïve pluripotency and differentiation markers (Fig. 2f and Extended Data Fig. 3d). To test if loser cells are developmentally delayed or advanced compared to control cells we projected our data onto a previously published diffusion map that includes epiblast cells from E5.5, E6.25 and E6.5 stage embryos^26^. We found that all epiblast cells, irrespective of the condition the embryos were cultured in (ie, DMSO or CI-treated) and of their losing state (ie, that they belonged to the winner, intermediate or loser cluster) mostly overlap with the E6.5 epiblast cells (Extended Data Fig. 3e-g). Cells from the loser cluster are slightly closer to the E6.25 stage than the winner and intermediate cells, as shown by their pseudo-time coordinate, but they remain far from the earlier E5.5 stage. This result combined with the higher expression of some differentiation markers observed in loser cells suggests that these cells are miss-patterned rather than developmentally delayed.

### Loser cells are characterised by defects in mitochondrial function

We next analysed using IPA the cellular pathways mis-regulated in loser epiblast cells and found that the top two pathways (mitochondrial dysfunction and oxidative phosphorylation) are related to mitochondrial function (Fig. 3a-b, Supplementary Table 1 and 2). For example, we found a down-regulation along the winner to loser trajectory of the mtDNA encoded *mt-Nd3* and *mt-Atp6*, of regulators of mitochondrial dynamics such as *Opa1,* as well as of genes involved in mitochondrial membrane and cristae organisation such as *Samm50* (Fig. 3c), suggesting that mitochondrial function is impaired in loser cells.

**Fig. 3.**
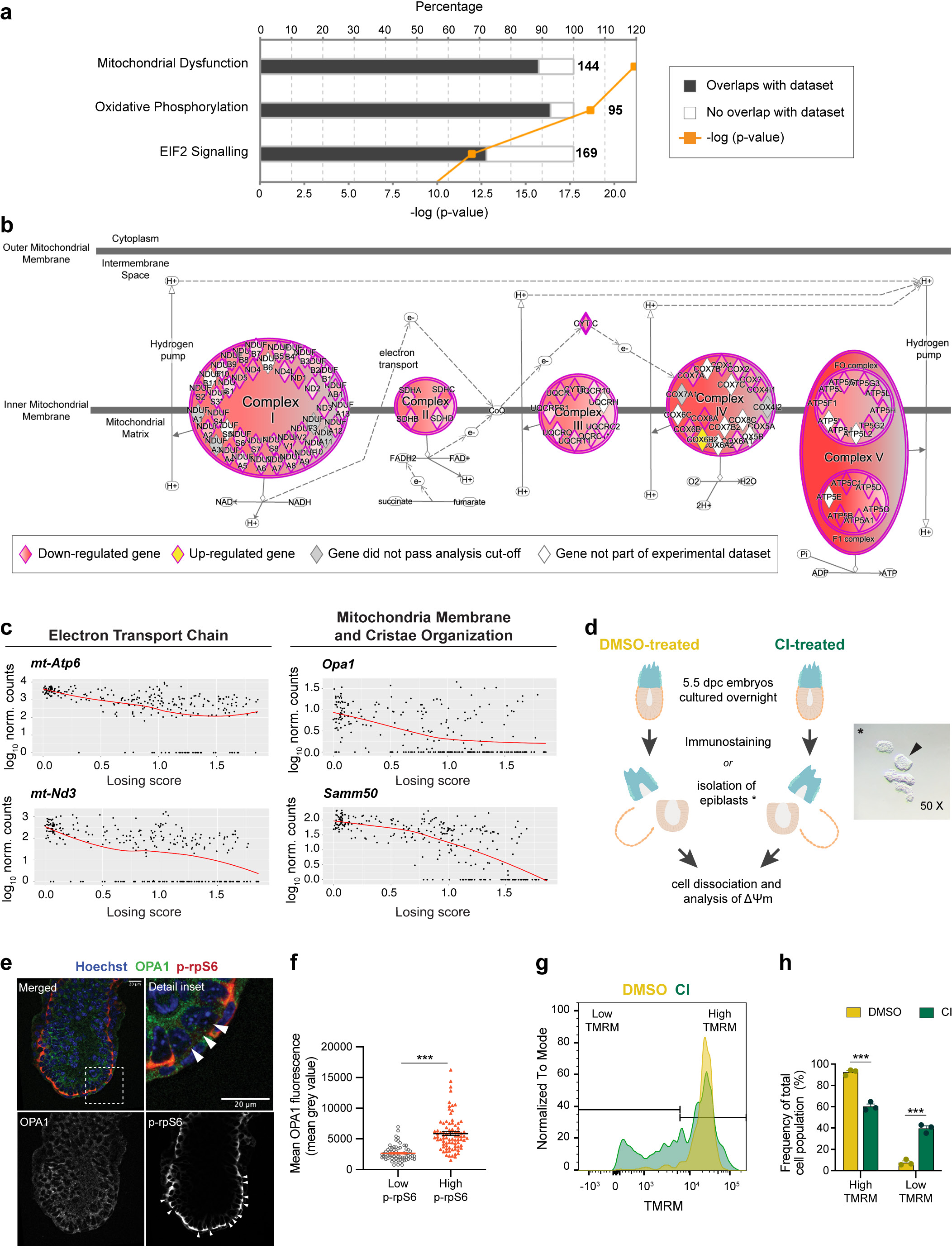
Cells eliminated during early mouse embryogenesis have mitochondrial defects. **a**, Top canonical pathways, identified by IPA, mis-regulated in loser cells in comparison to normal epiblast cells. The numbers at the end of each bar refer to total amount of genes involved in that pathway. The percentage refers to the number of genes found mis-regulated in loser cells relative to the number total genes within each pathway. Statistical significance calculated with Fisher’s exact test (p<0.05): Mitochondrial Dysfunction, -log_10_(p-value) = 21.1; Oxidative Phosphorylation, - log_10_(p-value) = 18.6; EIF2 signalling, - log_10_(p-value) = 11.9. **b**, Detail of changes in oxidative phosphorylation pathway identified in (**a**). Circular and oval shapes represent each of the ETC complexes (complexes I to V). Diamond shapes represent subunits of each ETC complex. Genes that are down-regulated in loser cells are coloured in shades of red. Darker shades correspond to lower values of FDR, which ranges from 1.25E-51 (for *Atp5b*) to 5.42E-03 (for *Ndufa11*). *Cox6b2,* coloured in yellow, was found to be up-regulated in loser cells (FDR=2.69E-13). Grey colour denotes genes that were not differentially expressed between loser and winner cells (FDR>0.01). White colour denotes genes from the Knowledge Base that were not tested (e.g., because they were not detected in our dataset). **c**, Expression levels of some mitochondrial genes as a function of cells’ losing score. *mt-Atp6*, mitochondrial DNA encoded ATP synthase membrane subunit 6; *mt-Nd3*, mitochondrial DNA encoded NADH dehydrogenase subunit 3; *Opa1*, optic atrophy 1; *Samm50*, sorting and assembly machinery component 50 homolog. **d**, Experimental design adopted to assess mitochondria function (OPA1 expression, by immunofluorescence or Δψm, given by TMRM fluorescence) in epiblast cells from embryos where cell death was allowed (DMSO-treated) or inhibited (CI-treated). * Micrograph of isolated epiblast (arrow) after embryo microdissection. **e**, Representative immunohistochemistry of OPA1 in E6.5 embryo where cell death was inhibited (CI-treated), quantified in (**f**). Loser cells are identified by low mTOR activation (low p-rpS6, arrowheads). Scale bar = 20 μm. **f**, Quantification of OPA1 fluorescence in normal epiblast cells and loser cells. N=6 embryos with a minimum of 8 cells analysed per condition. Statistical analysis performed by Mann-Whitney test. **g**, Representative histogram of flow cytometry analysis of TMRM probe, indicative of Δψm, in epiblast cells from embryos where cell death was allowed (DMSO-treated) or inhibited (CI-treated), quantified in (**h**). **h**, Frequency of epiblast cells with high or low TMRM fluorescence, according to range defined in (**g**) from embryos where cell competition was allowed (DMSO treated) or inhibited (CI-treated). Statistical analysis done by two-way ANOVA, followed by Holm-Sidak’s multiple comparisons test. N=3 independent experiments. Data shown as mean ± SEM.

A recent body of evidence has revealed that stress responses, such as the integrated stress response (ISR) or the closely related unfolded protein response (UPR), when triggered in cells with impaired mitochondrial function prompt a transcriptional program to restore cellular homeostasis^27–29^. We observed that loser epiblast cells displayed a characteristic UPR-ISR signature^30–33^ and key regulators of this response, such as *Atf4, Ddit3*, *Nrf2* and *Foxo3* were all up-regulated in these cells (Extended Data Fig. 4a-d). Similarly, *Sesn2*, a target of p53 that controls mTOR activity^34^, was also up-regulated in loser cells (Extended Data Fig. 4d). These findings support that loser epiblast cells present mitochondrial defects, leading to the activation of a stress response in an attempt to restore cellular homeostasis^35^.

To validate the significance of the observed mitochondrial defects, we did two things. First, we asked if the changes of mitochondrial regulators at the mRNA level are also reflected at the protein level. We observed that in CI-treated embryos, loser cells that persist and are marked by low mTOR activity^7^, also show significantly lower OPA1 levels (Fig. 3d-f). We also found that DMSO-treated embryos showed strong DDIT3 staining (an UPR-ISR marker) in the dying cells that accumulate in the proamniotic cavity, and that in CI-treated embryos, DDIT3 expression was up-regulated in a proportion of epiblast cells (Extended Data Fig. 4e-g). The second thing we did to validate the importance of the mitochondrial defects was to study in loser epiblast cells their mitochondrial membrane potential (Δψm), an indication of mitochondrial health. We observed that while the cells of DMSO-treated embryos showed a high Δψm that fell within a narrow range, in CI-treated embryos the proportion of cells with a low Δψm significantly increased (Fig. 3d and 3g-h). Together, these results suggest that loser epiblast cells have impaired mitochondrial activity that triggers a stress response.

### Mitochondrial dysfunction is common to different types of loser cells

To address if mitochondrial defects are a common feature of loser cells eliminated by cell competition, we analysed ESCs that are defective for BMP signalling (*Bmpr1a^-/-^)* and tetraploid cells (4n)^6^. We first carried out a mass spectrometry analysis using the Metabolon platform and found that metabolites and intermediates of the TCA cycle, such as malate, fumarate, glutamate and α-ketoglutarate are depleted in both *Bmpr1a^-/-^* and 4n ESCs in differentiation culture conditions (Fig. 4a). Next, we performed an extracellular flux Seahorse analysis of *Bmpr1a^-/-^* ESCs to measure their glycolytic and oxidative phosphorylation (OXPHOS) rates. We observed that when these cells are maintained in pluripotency culture conditions that are not permissive for cell competition^6^, they showed a similar glycolytic activity but a higher OXPHOS rate than control cells (Extended Data Fig. 5a-b). In contrast, when *Bmpr1a^-/-^* cells are induced to differentiate, this phenotype is reversed, with mutant cells showing lower ATP generated through OXPHOS and a higher glycolytic capacity than controls (Fig. 4b-e and Extended Data Fig. 5c-d). This suggests that upon differentiation *Bmpr1a^-/-^* cells are unable to sustain proper OXPHOS activity.

**Fig. 4.**
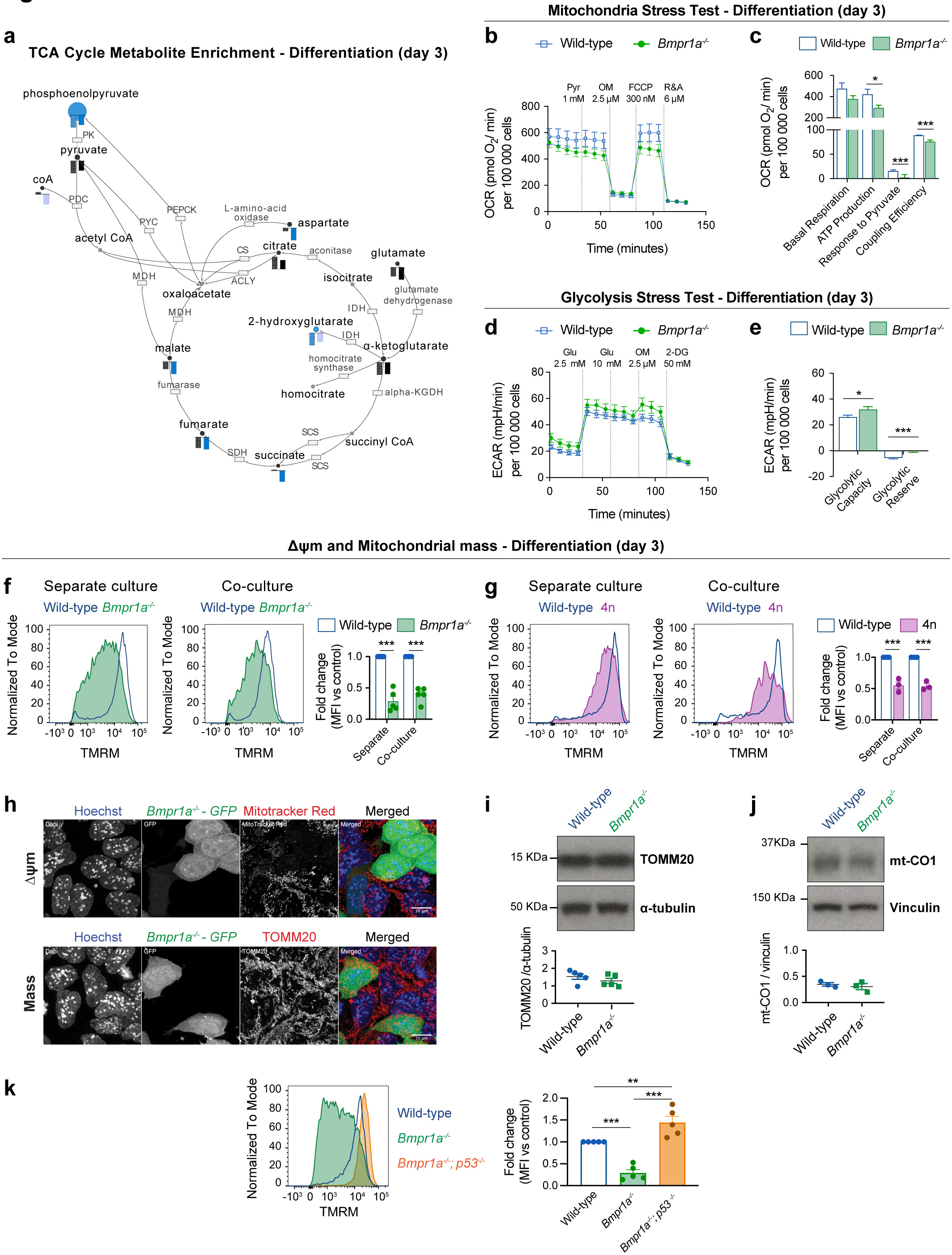
Mitochondrial defects are a common feature of cells eliminated by cell competition. **a**, Metabolic enrichment analysis of the TCA cycle and intermediate metabolites obtained using Metabolon platform for defective cells (*Bmpr1a^-/-^*, left bar and 4n, right bar), in comparison to wild-type cells during differentiation. Bars indicate compound levels relative to wild-type cells. Blue bars indicate compounds that are significantly depleted (p<0.05) and light blue bars indicate compounds that are almost significantly depleted (0.05≤p≤0.1). Black bars indicate compounds that are depleted although not statistically significant in comparison to the levels found in wild-type cells. The enzymes on the pathway are represented as boxes and labelled by their canonical names. **b-e**, Metabolic flux analysis of wild-type and BMP-defective cells during differentiating conditions. Data obtained with a minimum of 3 independent experiments, with 5 replicates per cell type in each assay. Statistical analysis done with Mann-Whitney test. Analysis of oxygen consumption rate (OCR) as a measure of mitochondria function (mitochondria stress test) (**b**). Detail of metabolic parameters found changed from the analysis of the mitochondria stress test (**c**). Analysis of extracellular acidification rate (ECAR) as a measure of glycolytic function (glycolysis stress test) (**d**). Detail of metabolic parameters found changed from the analysis of the glycolysis stress test (**e**). **f**-**g**, Analysis of mitochondrial membrane potential (Δψm) in defective mESCs undergoing differentiation in separate or co-culture conditions. Representative histograms of TMRM fluorescence and quantification for wild-type and *Bmpr1a^-/-^* (**f**) and wild-type and 4n (**g**). Statistical analysis done by two-way ANOVA, followed by Holm-Sidak’s multiple comparisons test. **h**, Representative micrographs of wild-type and *Bmpr1a^-/-^* cells co-cultured during differentiation and stained for a reporter of Δψm (MitoTracker Red, top panel) or mitochondria mass (TOMM20, bottom panel). Nuclei are stained with Hoechst. Scale bar = 10 μm. **i**-**j**, Western blot analysis of mitochondria mass markers TOMM20 (**i**) and mt-CO1 (**j**) for wild-type and *Bmpr1a^-/-^* cells during differentiation. Statistical analysis done with Mann-Whitney test (**i**) or unpaired t-test (**j**). **k**, Analysis of mitochondrial membrane potential (Δψm) for wild-type, *Bmpr1a^-/-^* and *Bmpr1a^-/-^*;*p53^-/-^* cells during differentiation. Representative histogram of TMRM fluorescence and quantification. Statistical analysis done by one-way ANOVA, followed by Holm-Sidak’s multiple comparisons test. Data shown as mean ± SEM of a minimum of 3 independent experiments.

To further test the possibility that defective ESCs have impaired mitochondrial function, we assessed their Δψm. We found that whilst *Bmpr1a^-/-^* and 4n cells had a similar Δψm to control cells in pluripotency conditions (Extended Data Fig. 5e-f), upon differentiation both these cell types presented a loss of Δψm, irrespective of whether they were separate or co-cultured with wild-type cells (Fig. 4f-g). This reduction in Δψm is not due to excessive mitochondrial reactive oxygen species (ROS) production or to a lower mitochondrial mass within mutant cells since, as for example, *Bmpr1a^-/-^* cells have lower ROS levels and similar TOMM20 and mt-CO1 expression as control cells (Fig. 4h-j and Extended Data Fig. 5g). The fact that the loss of Δψm and lower OXPHOS activity can be observed even when loser cells are cultured separately, suggests that the mitochondrial dysfunction phenotype is an inherent property of loser cells and not a response to them being out-competed. These results also indicate that the mitochondrial defects are directly linked to the emergence of the loser status: in conditions that are not permissive for cell competition (pluripotency) mutant cells do not show defective mitochondrial function, but when they are switched to differentiation conditions that allow for cell competition, they display impaired mitochondrial function.

To further explore the relationship between mitochondrial activity and the competitive ability of the cell, we analysed the Δψm of BMP defective cells that are null for p53 (*Bmpr1a^-/-^; p53^-/-^* ESCs), as these are not eliminated by wild-type cells^7^. Remarkably, we observed that mutating *p53* in *Bmpr1a^-/-^* cells not only rescues the loss of Δψm of these cells, but also causes hyperpolarisation of their mitochondria (Fig. 4k). These results not only suggest a role for P53 in regulating mitochondrial activity of ESCs, but also strongly support a pivotal role for mitochondrial activity in cell competition.

### Impaired mitochondrial function is sufficient to trigger cell competition

The mitochondrial defects observed in loser cells led us to ask if disrupting mitochondrial activity alone is sufficient to trigger cell competition. During the onset of differentiation, mitochondrial shape changes substantially. In pluripotent cells mitochondria have a round and fragmented shape, but upon differentiation they fuse and become elongated, forming complex networks^10^.

Given that this change in shape correlates with when cell competition occurs, we tested if disrupting mitochondrial dynamics is sufficient to induce cell competition. MFN1 and MFN2 regulate mitochondrial fusion and DRP1 controls their fission^36–38^. We generated *Mfn2^-/-^* ESCs, that have enlarged globular mitochondria, and *Drp1^-/-^* ESCs, that show hyper-elongated mitochondria (Fig. 5a). We observed that *Mfn2^-/-^* ESCs displayed very poor growth upon differentiation (data not shown). For this reason, we tested their competitive ability in pluripotency conditions, that we have previously found not to induce the out-competition of *Bmpr1a^-/-^* or 4n cells^6^. Interestingly, we found that although *Mfn2^-/-^* cells grow similarly to wild-type cells in separate cultures, they were out-competed in co-culture (Fig. 5b). Analysis of the *Drp1* mutant cells showed that although they did not grow significantly slower than wild-type cells when cultured separately in differentiation inducing conditions, they were out-competed by wild-type cells in co-culture (Fig. 5c). The observation that disrupting mitochondrial dynamics can induce cell competition even in pluripotency culture conditions, suggests that mitochondrial activity is a dominant parameter determining the competitive ability of the cell.

To establish how disruption of mitochondrial fusion and fission affects mitochondrial performance we compared the Δψm, respiration rates and mitochondrial ATP production of *Mfn2^-/-^* and *Drp1^-/-^* ESCs to that of wild-type cells (Fig. 5d-g). We found that whilst *Mfn2^-/-^* and *Drp1^-/-^* ESCs had lower Δψm than control cells (Fig. 5d,f), *Mfn2^-/-^* ESCs had lower maximal respiration rates but similar basal respiration and ATP production to controls and *Drp1^-/-^* ESCs showed similar respiration and ATP production to controls (Fig. 5e,g). This suggests that ATP production or respiration rates alone do not determine the relative competitive ability of ESCs.

**Fig. 5.**
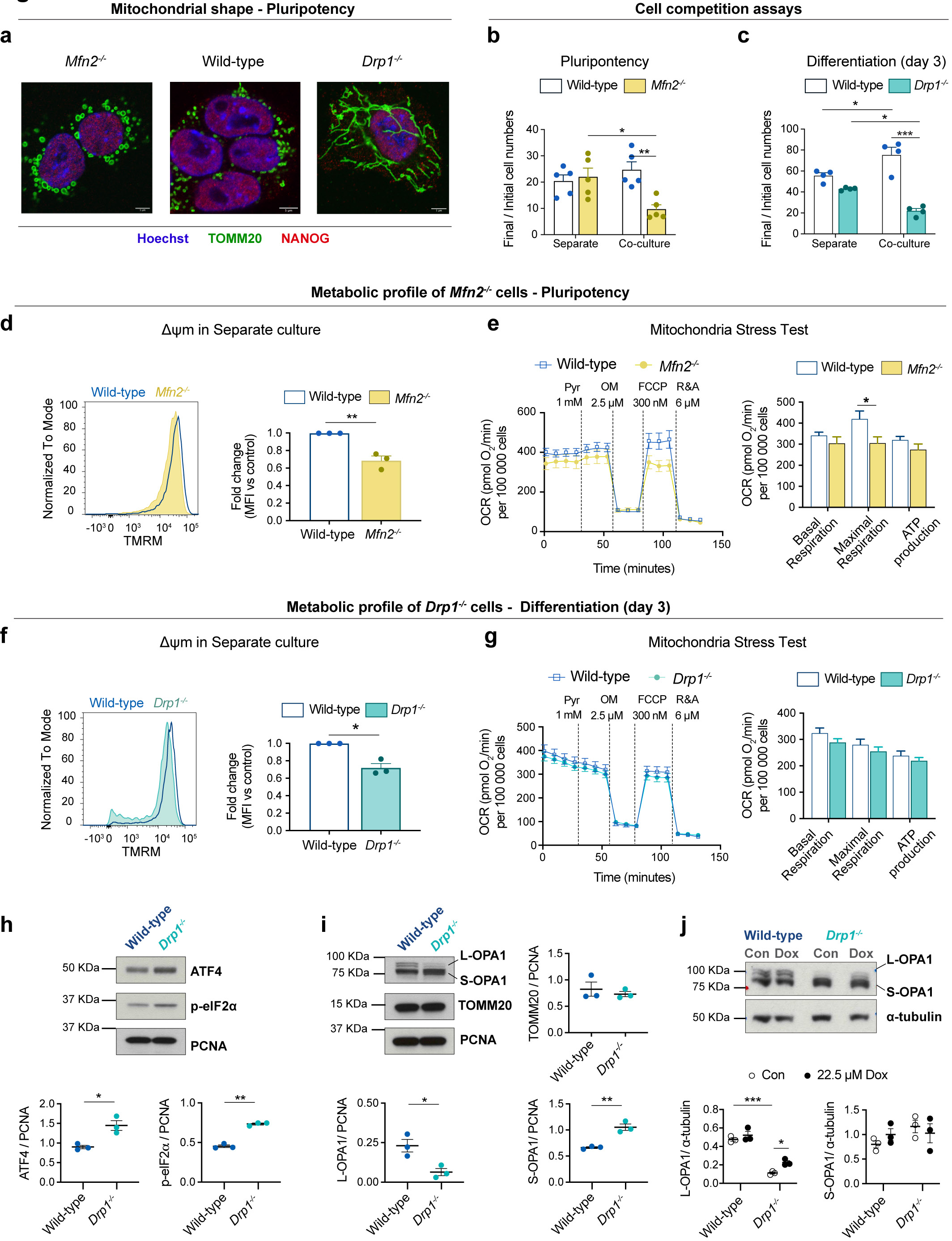
Manipulating mitochondria biology is sufficient to trigger cell competition. **a**, Representative micrographs of wild-type*, Mfn2^-/-^* and *Drp1^-/-^* mESCs showing alterations in mitochondrial morphology in mutant cells. TOMM20 was used as a mitochondrial marker and NANOG as a pluripotency marker. Nuclei are stained with Hoechst. Scale bar = 5 μm. **b**,**c**, Cell competition assays between wild-type mESCs and cells with altered morphology, *Mfn2^-/-^* during pluripotency (**b**) and *Drp1^-/-^* during differentiation (**c**). The ratio of final/initial cell numbers in separate or co-culture is shown. Statistical analysis done with two-way ANOVA, followed by Holm-Sidak’s multiple comparisons test. **d-j**, Metabolic profile of *Mfn2^-/-^* and *Drp1^-/-^* mESCs. Analysis of mitochondrial membrane potential (Δψm) for wild-type and *Mfn2^-/-^* cultured separately during pluripotency (**d**) and for wild-type and *Drp1^-/-^* mESCs *^-/-^* after 3 days of differentiation in separate culture (**f**). Data was obtained from 3 independent experiments and statistical testing done with one sample t-test. Metabolic flux analysis of wild-type and *Mfn2^-/-^* mESCs cultured separately during pluripotency (**e**) and for wild-type and *Drp1^-/-^* after 3 days of differentiation in separate cultures (**g**). Data was collected from 3 independent experiments, with 5 replicates per cell type in each assay, and statistical testing done with Mann-Whitney test. **h-j**, Western blot analysis of markers of UPR and mitochondrial markers in wild-type and *Drp1^-/-^* after 3 days of differentiation in separate culture. Cells were treated with doxycycline (Dox, 22.5 μM) or vehicle (Con) from day 1 of differentiation and samples were collected on day 3 (**j**). Statistical analysis was done with an unpaired t-test (h-i) or two-way ANOVA followed by Holm-Sidak’s multiple comparisons test (j). Data shown as mean ± SEM of a minimum of 3 independent experiments. p-eIF2ɑ, phosphorylated eukaryotic initiation factor 2ɑ.

Besides mitochondrial dysfunction, another prominent signature of loser cells found *in vivo* was the UPR/ISR (Ext. Data Fig. 4). Since the loss of *Drp1* has been associated with activation of the UPR^39–41^, we investigated if the *Drp1^-/-^* loser cells also showed evidence for the activation of the UPR/ISR. We observed that *Drp1^-/-^* cells show higher expression of ATF4 and p-eIF2ɑ than wild-type counterparts, which is indicative of UPR/ISR activation (Fig. 5h)^39–41^. Another feature previously described upon loss of *Drp1* is the proteolytic cleavage of OPA1, where short isoforms (S-OPA1) are accumulated in detriment of the long isoforms (L-OPA1)^39^. When we analysed the expression of OPA1 in wild-type and *Drp1^-/-^* cells we observed that while wild-type cells retain L-OPA1 expression, loser cells predominantly express the S-OPA1 isoforms and display almost no expression of L-OPA1 (Fig. 5i). This defect has been associated with mito-ribosomal stalling, a phenotype that can be replicated by treating cells with actinonin (Extended Data Fig. 6)^42^. To test if the shift in isoform expression observed in *Drp1^-/-^* ESCs is due to aberrant mitochondrial translation we treated cells with doxycycline, that inhibits this translation^43^, and observed that this was sufficient to partially rescue L-OPA1 expression (Fig. 5j). This rescue together with the evidence for UPR/ISR expression suggest that *Drp1^-/-^* cells display defects in mitochondrial translation.

### Loser epiblast cells accumulate mtDNA mutations

There is strong evidence for selection against aberrant mitochondrial function induced by deleterious mtDNA mutations in mammals^21, 44–47^. Given that we observe that cell competition selects against cells with impaired mitochondrial function, we asked if cell competition could be reducing mtDNA heteroplasmy (frequency of different mtDNA variants) during mouse development. It has been recently shown that scRNA-seq can be used to reliably identify mtDNA variants, although with a lower statistical power compared to more direct approaches, like mtDNA sequencing^48^. We therefore tested if mtDNA heteroplasmy is present in our scRNA-seq data and whether this correlates with the losing score of a cell. Our analysis revealed that the frequency of specific mtDNA polymorphisms increased with the losing score of epiblast cells (Fig. 6a), and such mtDNA changes occurred within *mt-Rnr1* and *mt-Rnr2* (Fig. 6b-h and Extended Data Fig. 7a-e). Moreover, these changes were not dependent on the litter from which the embryo came from (Extended Data Fig. 7f-k). The mutations we detected in *mt-Rnr1* and *mt-Rnr2* strongly co-occurred in the same cell, with those closest together having the highest probability of co-existing (Fig. 6i and Extended Data Fig. 7l). This is suggestive of mtDNA replication errors that could be ‘scarring’ the mtDNA, disrupting the function of *mt-Rnr1* (12S rRNA) and *mt-Rnr2* (16S rRNA) and causing the loser phenotype. Importantly, the presence of these specific mtDNA mutations in the loser cells suggests that cell competition is contributing to the elimination of deleterious mtDNA mutations during early mouse development.

**Fig. 6.**
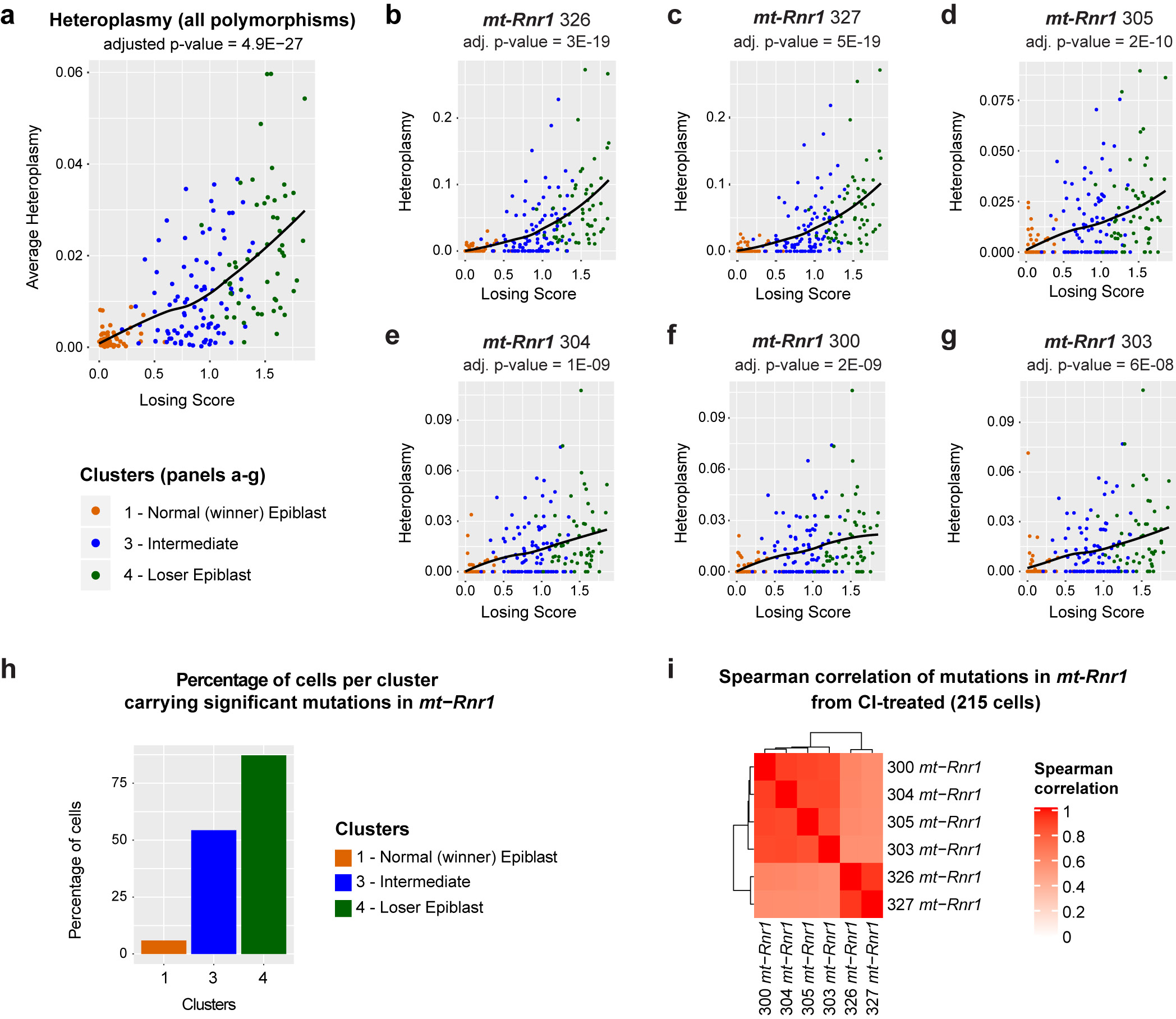
Intermediate and loser epiblast cells accumulate polymorphisms in mtDNA sequence. **a-g**, mtDNA heteroplasmy in epiblast cells from CI-treated embryos. Average heteroplasmy (considering all eleven polymorphisms that have a statistically significant dependence on the losing score; see Methods) as a function of cells’ losing scores. The p-value was computed with a generalized linear model (**a**). mtDNA heteroplasmy for six positions within the *mt-Rnr1* gene (**b-g**). The heteroplasmy at all of these positions as well as the average heteroplasmy increase with the cells’ losing scores in a statistically significant way (the adjusted p-value estimated via a generalized linear model is indicated at the top of each plot). **h**, The barplot indicates the fraction of epiblast cells in each of the cluster indicated on the x-axis (winner, intermediate, loser) that carry a mean heteroplasmy (computed on the six positions within the *mt-Rnr1* indicated in the panels **b**-**g**) greater than 0.01. This shows that the level of mtDNA heteroplasmy in *mt-Rnr1* is strongly associated with the loser status of the cells, since ∼55% and ∼87% of cells in the intermediate and the loser clusters, respectively, have heteroplasmic sequences in this gene compared to only ∼5% of cells in the winner cluster. **i**, Spearman’s correlation coefficient between the mtDNA heteroplasmy at the six positions shown in panels (**b**-**g**).

### Changes in mtDNA sequence can determine the competitive ability of a cell

To explore this possibility further, we analysed if alterations in mtDNA can induce cell competition by testing the competitive ability of ESCs with non-pathological differences in mtDNA sequence. For this we compared the relative competitive ability of ESCs that shared the same nuclear genome background but differed in their mitochondrial genomes by a small number of non-pathological sequence changes. We derived ESCs from hybrid mouse strains that we had previously engineered to have a common nuclear C57BL/6N background, but mtDNAs from different wild-caught mice^16^. Each wild-derived mtDNA variant (or haplotype) contains a specific number of single nucleotide polymorphisms (SNPs) that lead to a small number of amino acid changes when compared to the C57BL/6N mtDNA haplotype. Furthermore, these haplotypes (BG, HB and ST) can be ranked according to their genetic distance from the C57BL/6N mtDNA (Fig. 7a and Extended Data Fig. 8a). Characterization of the isolated ESCs revealed that they have a range of heteroplasmy (mix of wild-derived and C57BL/6N mtDNAs) that is stable over several passages (Extended Data Fig. 8b). Importantly, these different mtDNA haplotypes and different levels of heteroplasmy do not alter cell size, cell granularity, mitochondrial mass or mitochondrial dynamics, nor do they substantially impact the cell’s Δψm (Extended Data Fig. 8c-f).

**Fig. 7.**
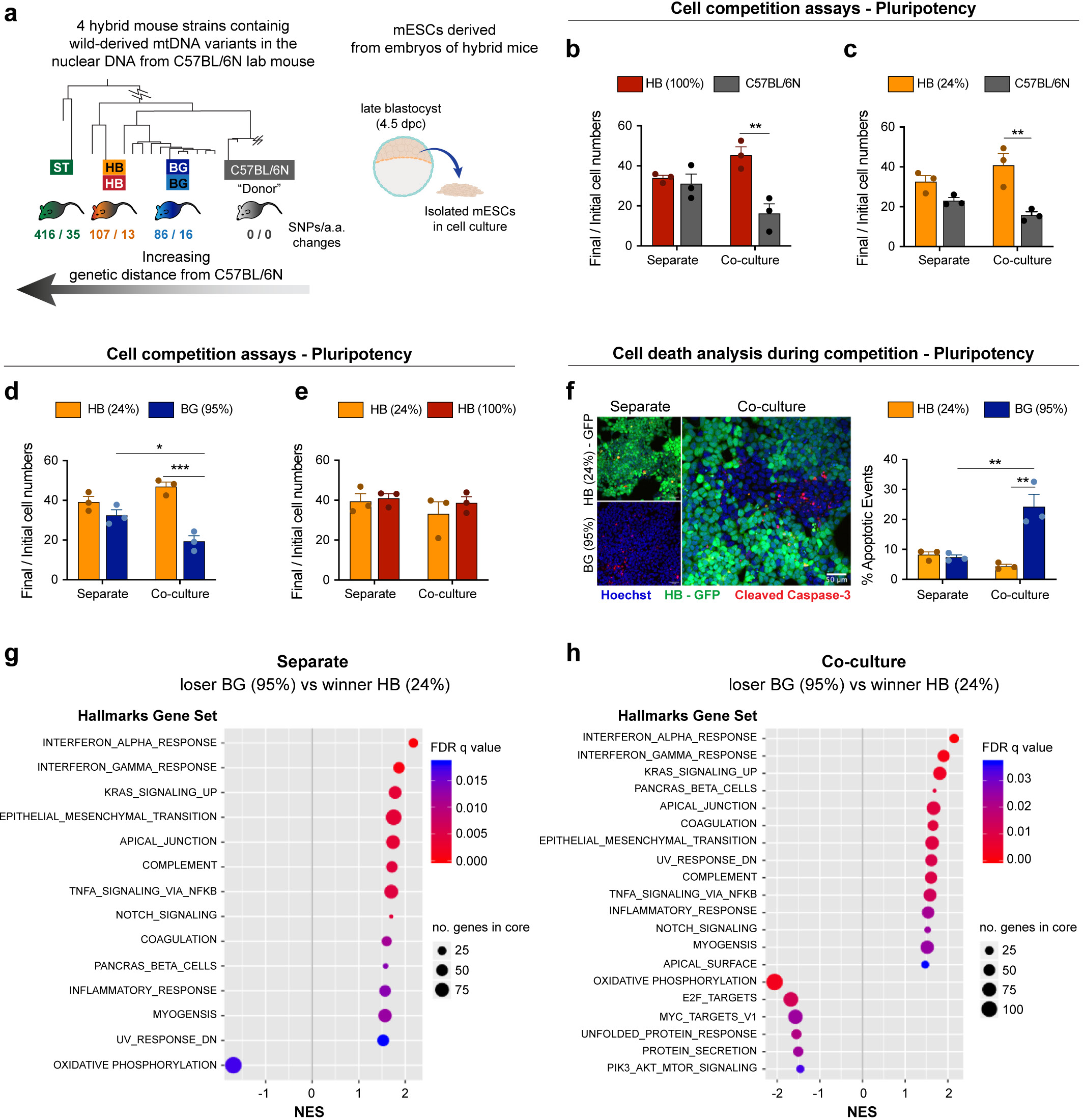
Changes in mtDNA sequence can determine the competitive ability of a cell. **a,** Derivation of mESCs from hybrid mouse strains, generated elsewhere by Burgstaller and colleagues. Neighbour-Joining Phylogenetic Analysis of mtDNA from wild and C57BL/6N mouse strains, that were used to generate hybrid mice (adapted from^16^), illustrates the genetic distance of the mtDNA from wild mouse strains to the C57BL/6N lab mouse. The number of single nucleotide polymorphisms and amino acid changes (SNPs/ a.a. changes) from wild to lab mouse strain is shown. mESCs were derived from embryos of hybrid mice, containing the nuclear background of a C57BL/6N lab mouse and mtDNA from three possible wild-derived strains (BG, HB or ST). **b-e,** Cell competition assays between cells derived from the embryos of hybrid mice performed in pluripotency maintenance conditions. The ratio of final/initial cell numbers in separate or co-culture is shown. Statistical analysis done with two-way ANOVA, followed by Holm-Sidak’s multiple comparisons test. **f,** Representative micrographs of cleaved caspase-3 staining and quantification of the percentage of apoptotic events in winners HB(24%) and loser BG(95%) mESCs maintained pluripotent and cultured in separate or co-culture conditions. Statistical analysis done with two-way ANOVA, followed by Holm-Sidak’s multiple comparisons test. **g-h,** Gene set enrichment analysis of differentially expressed genes from bulk RNA seq. in loser BG (95%) compared to winner HB (24%) mESCs maintained pluripotent and cultured in separate (**g**) or co-culture conditions (**h**). Gene sets that show positive normalized enrichment scores (NES) are enriched in loser cells, while gene sets that show negative NES are depleted in loser cells. Data in panels (**b**-**f**) shown as mean ± SEM of a minimum of 3 independent experiments.

When we tested the competitive ability of these ESCs with different mtDNA content, in pluripotency culture conditions, we observed that cells carrying the mtDNAs that are most distant from the C57BL/6N mtDNA, such as the HB(100%), the HB(24%) and the ST(46%) ESCs could all out-compete the C57BL/6N line (Fig. 7b-c and Extended Data Fig. 8g). Similarly, when we tested the HB(24%) line against the BG(99%) or the BG(95%) lines (that have mtDNAs more closely related to the C57BL/6N mtDNA), we found that cells with the HB haplotype could also out-compete these ESCs (Fig. 7d and Extended Data Fig. 8h). In contrast, we observed that the HB(24%) ESCs were unable to out-compete either their homoplasmic counterparts, HB(100%), or the ST(46%) cells that carry the most distant mtDNA variant from C57BL/6N (Fig. 7e and Extended Data Fig. 8i). These results tell us three things. First, that non-pathological differences in mtDNA sequence can trigger cell competition. Second, that a competitive advantage can be conferred by only a small proportion of mtDNA content, as indicated by our finding that HB(24%) behave as winners. Finally, these findings suggest that the phylogenetic proximity between mtDNA variants can potentially determine their competitive cell fitness.

To characterise the mode of competition between cells with different mtDNA we focussed on the HB(24%) and the BG(95%) ESCs. Analysis of these cell lines revealed that specifically when co-cultured, the BG(95%) cells display high levels of apoptosis (Fig. 7f), indicating that their out-competition is through their elimination. To gain further insight we performed bulk RNA-seq of these cells in separate and co-culture conditions (Extended Data Fig. 8j) and analysed the differentially expressed genes by gene-set enrichment analysis (GSEA). We found that in separate culture the most notable features that distinguish BG(95%) from HB(24%) cells were a down-regulation of genes involved in oxidative phosphorylation and an up-regulation of those associated with cytokine activity (Fig. 8g). Interestingly, in the co-culture condition, in addition to these signatures, BG(95%) cells revealed a down-regulation in signature markers of MYC activity and mTOR signalling (Fig. 7h), whose downregulation are known read-outs of a loser status during cell competition in the embryo^5–7^ (Fig. 2c).

**Fig. 8.**
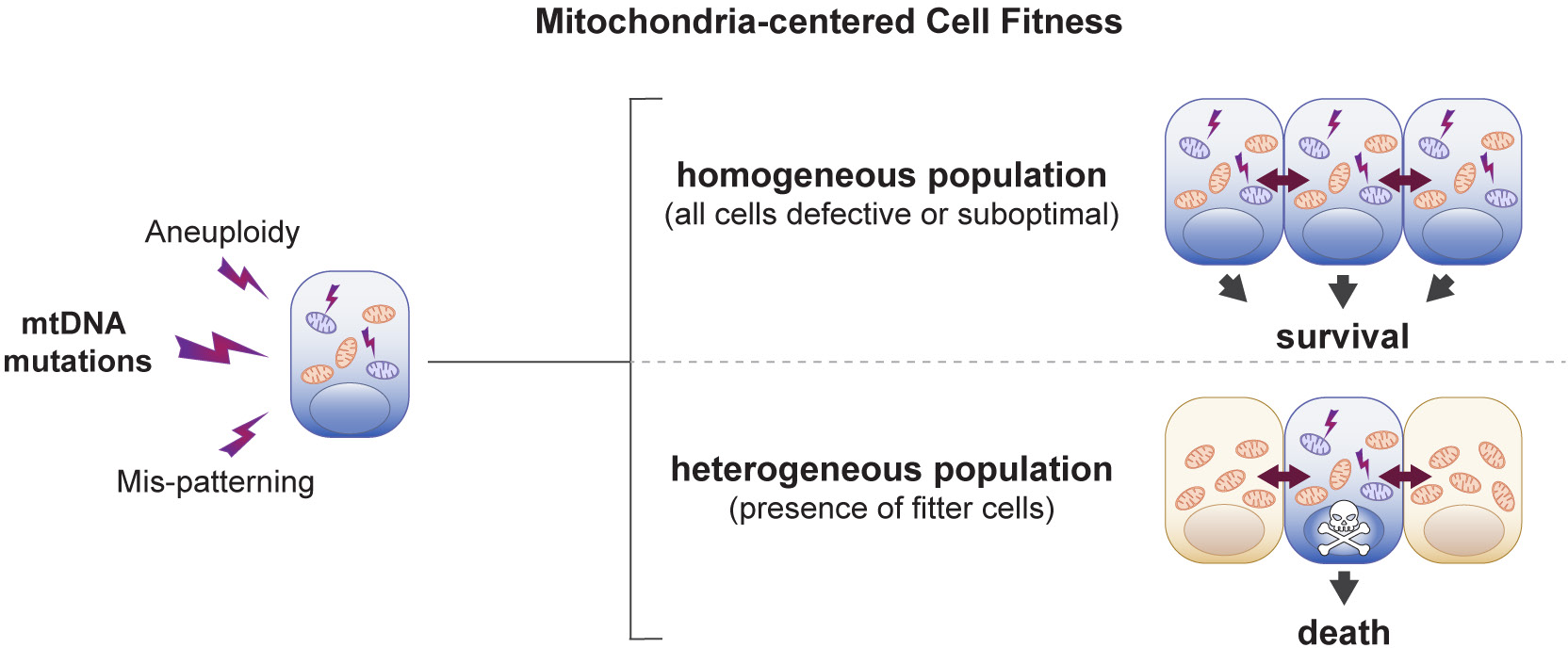
Model of cell competition. Summary of the main findings of the study. A range of cellular defects, such as aneuploidy, mis-patterning or mtDNA mutations cause alterations in mitochondria function, which affect the relative fitness of cells. The cells with suboptimal mitochondrial activity survive in a homogeneous population but are eliminated by cell competition in the presence of fitter cells.

To test if the down-regulation of genes involved in oxidative phosphorylation was also reflected at the functional level we compared oxygen consumption rates and mitochondrial ATP generation in HB(100%), HB(24%), BG(95%) and C57Bl/6N ESCs. We find that the winner cells HB(100%) and HB(24%) have higher basal respiration, higher maximal respiration and higher mitochondrial ATP production than the loser BG(95%) and C57BL/6N ESCs (Extended Data Fig. 9). These data indicate that the mtDNA differences that exist between winner and loser cells are sufficient to affect their mitochondrial performance and this ultimately determines their competitive ability. However, the observation that differentiating *Drp1^-/-^* ESCs are eliminated by cell competition but do not show differences in respiration rates or mitochondrial ATP production (Fig. 5b,e), suggests that respiration or ATP production rates alone are unlikely to be the mitochondrial parameters that control competitive cell fitness.

The finding that the genes down-regulated in BG(95%) cells when co-cultured with HB(24%) cells fell under functional categories relating to mitochondrial function (Extended Data Fig. 10a) led us to analyse the degree of overlap between these genes and the genes differentially expressed along the winner-to-loser trajectory in the embryo. We observed a significant overlap in mis-regulated genes (Extended Data Fig. 10b), as well as in the functional components that these genes can be categorised into (Extended Data Fig. 10c). This further highlights the importance of relative mitochondrial activity for determining the competitive ability of embryonic cells.

## Discussion

The emerging role of cell competition as a regulator of cell fitness in a wide range of cellular contexts, from the developing embryo to the ageing tissue^1–3^, has highlighted the importance of understanding what cell types are normally eliminated by this process. With the aim of understanding this question, we have analysed the transcriptional identity of the cells eliminated in the early mouse embryo. We have found not only that they present a cell competition signature but also that they are marked by mtDNA mutations and impaired mitochondrial function. Starting from these results, we leveraged *in vitro* models of cell competition to show that: (i) mitochondrial function is impaired in loser cells eliminated by cell competition, and (ii) differences in mitochondrial activity are sufficient to trigger cell competition in ESCs. Overall, this points to mitochondrial performance as a key determinant of the competitive ability of cells during early mammalian embryonic development. One implication of our findings is that a range of different types of defects, such as mis-patterning, karyotypic abnormalities or mtDNA mutations, all lead to dysfunctional mitochondria at the onset of differentiation and that ultimately it is their impaired mitochondrial function that triggers cell competition, inducing their elimination (Fig. 8).

Embryos are exposed to different microenvironments in vivo and when cultured ex-vivo. Similarly, ESCs also experience a different micro-environment to epiblast cells in the embryo. These different micro-environments could potentially affect the selective pressure and hence the transcriptional signature of loser cells. However, there are two reasons why we think that the loser cell signatures we identify here are conserved across systems. First, the transcriptional profile of our epiblast cells from cultured embryos is very similar to that of epiblast cells from freshly isolated embryos (see Extended Data Figure 3e-g). Second, the loser signature identified here is enriched for targets of P53 and depleted for mTOR and c-MYC targets. Given that these are regulators of cell competition identified by us and others in the embryo and in ESCs^5–7^, it suggests that the same pathways are inducing loser cell elimination in in vivo, ex-vivo and in ESC models of cell competition.

It is well known that the successful development of the embryo can be influenced by the quality of its mitochondrial pool^10^. Moreover, divergence from normal mitochondrial function during embryogenesis is either lethal or can lead to the development of mitochondrial disorders^49^.

Deleterious mtDNA mutations are a common cause of mitochondrial diseases and during development selection against mutant mtDNA has been described to occur through at least two mechanisms: the bottleneck effect and the intra-cellular purifying selection. The bottleneck effect is associated specifically with the unequal segregation of mtDNAs during primordial germ cell specification, for example as seen in the human embryo^50^. In contrast to this, purifying selection, as the name implies, allows for selection against deleterious mtDNAs and has been proposed to take place both during development and post-natal life^51^. Importantly, purifying selection has been found to occur at the molecule and organelle level, as well as at the cellular level^52^. Our findings indicate that purifying selection can occur not only at the intra-cellular level but also inter-cellularly (cell non-autonomously). We show that epiblast cells are able to sense their relative mitochondrial activity and that those cells with mtDNA mutations, lower or aberrant mitochondrial function are eliminated. By selecting those cells with the most favourable mitochondrial performance, cell competition would not only prevent cells with mitochondrial defects from contributing to the germline or future embryo, but also ensure optimisation of the bioenergetic performance of the epiblast, therefore contributing to the synchronization of growth during early development.

Cell competition has been studied in a variety of organisms, from *Drosophila* to mammals, and it is likely that multiple different mechanisms fall under its broad umbrella^1–3^. In spite of this, there is considerable interest in understanding if there could be any common feature in at least some of the contexts where cell competition has been described. The first demonstration of cell competition in *Drosophila* was made by inducing clones carrying mutations in the ribosomal gene *Minute*^4^ and this has become one of the primary models to study this process. Our finding that during normal early mouse development cell competition eliminates cells carrying mutations in *mt-Rnr1* and *mt-Rnr2*, demonstrates that in the physiological context mutations in ribosomal genes also trigger cell competition. Furthermore, our observation that mis-patterned and karyotypically abnormal cells show impaired mitochondrial activity indicates that during early mouse development different types of defects impair mitochondrial function and trigger cell competition. Interestingly, mtDNA genes are amongst the top mis-regulated factors identified during cell competition in the mouse skin^53^. In the *Drosophila* wing disc oxidative stress, a general consequence of dysfunctional mitochondria, underlies the out-competition of *Minute* and *Mah-jong* mutant cells^54^. Similarly, in MDCK cells a loss of Δψm occurs during the out-competition of RasV12 mutant cells and is key for their extrusion^55^. These observations raise the possibility that differences in mitochondrial activity may be a key determinant of competitive cell fitness in a wide range of systems. Unravelling what mitochondrial features can lead to cellular differences that can be sensed between cells during cell competition and if these are conserved in human systems will be key not only for understanding this process, but also to open up the possibility for future therapeutic avenues in the diagnosis or prevention of mitochondrial diseases.

## Acknowledgments

We would like to thank Stephen Rothery for guidance and advice with confocal microscopy. The Facility for Imaging by Light Microscopy (FILM) at Imperial College London is part-supported by funding from the Wellcome Trust (grant 104931/Z/14/Z) and BBSRC (grant BB/L015129/1). We thank James Elliot and Bhavik Patel from the LMS/NIHR Imperial Biomedical Research Centre Flow Cytometry Facility for support. We are thankful to George Chennell and Alessandro Sardini for guidance and support with Seahorse experiments. We also want to acknowledge Thomas Kolbe from University of Veterinary Medicine (Vienna) for isolating embryos from hybrid mice strains from which mESCs were derived. Research in Tristan Rodriguez lab was supported by the MRC project grant (MR/N009371/1) and by the British Heart Foundation centre for research excellence. Work in the Scialdone lab is funded by the Helmholtz Association. Ana Lima was funded by a BHF centre of excellence PhD studentship. Shankar Srinivas was funded through Wellcome awards 103788/Z/14/Z and 108438/Z/15/Z.

## Author Contributions

A.L. performed most of the experimental wet lab work. J.B. and A.L. derived heteroplasmic mESC lines. J.B. performed heteroplasmy measurements in heteroplasmic mESCs. B.P. generated *Mfn2^-/-^* and *Drp1^-/-^* mESCs and J.M.S did characterisation of mitochondria shape and pluripotency status. S.P.M participated in the metabolic characterisation of *Drp1^-/-^* cells. D.H. performed embryo dissections, treatments and cell dissociation prior to scRNA-seq experiments.

G.L. did the bioinformatic analysis of scRNA-seq data. E.M., N.J. and A.G. participated in the analysis of mitochondrial DNA heteroplasmy. A.D.G. performed the metabolomic studies using Metabolon platform and participated in embryo dissections and immunohistochemistry stainings for validation of results obtained by scRNA-seq. M.D, and M.K. performed the bioinformatic analysis of bulk RNA-seq experiments. N.J., S.S. and D.C. participated in the design of experimental work and analysis of results. A.L., G.L., A.S and T.R. interpreted results and wrote the paper. T.R. and A.S. directed and designed the research.

## Competing Interests

The authors declare no competing interests.

## Figure titles and legends

**Extended Data Fig. 1.**
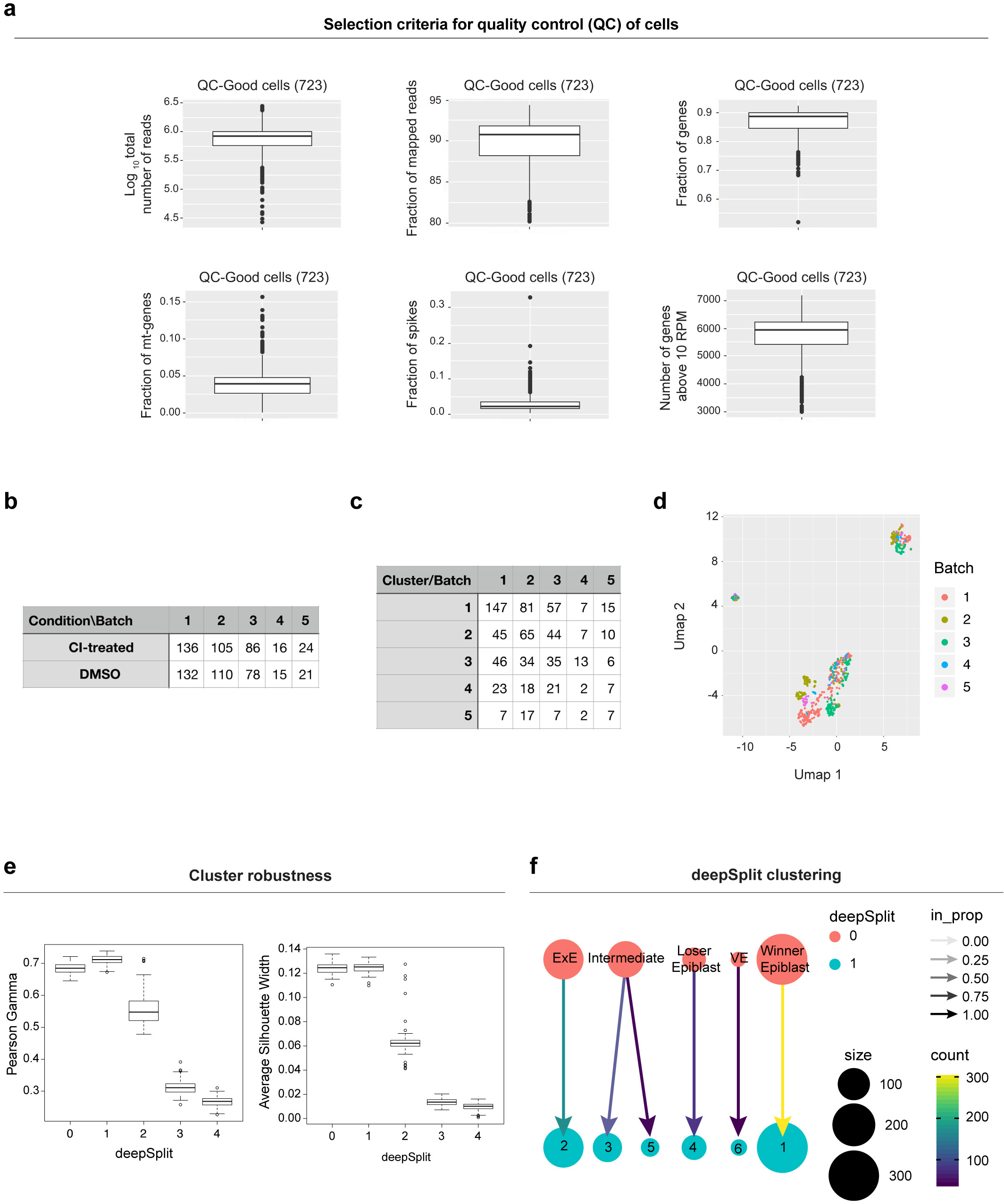
Quality controls of scRNA-seq and clustering robustness analysis. **a,** Selection criteria for quality control (QC) of all cells. A total of 723 passed the quality control (723 good quality cells) and were considered for downstream analysis. All these parameters were computed for each cell. Log_10_ total number of reads (top left): log_10_ of the sum of the number of reads that were processed in every cell; Fraction of mapped reads (top central): number of reads that are confidentially mapped to the reference genome divided by total number of reads that were processed for each cell. This number is automatically provided by Salmon v0.8.2; Fraction of genes (top right): number of reads mapped to endogenous genes divided by the total sum of reads that were processed; Fraction of mt-genes (bottom left): number of reads mapped to mitochondrial genes divided by the total sum of reads that were processed; Fraction of spikes (bottom central): number of reads mapped to ERCC spike-ins divided by the total sum of reads that were processed; Number of genes above 10 RPM (bottom right): number of genes with expression level above 10 reads per million. **b,** Number of good quality cells in each condition (rows) and batch (columns). **c,** Number of good quality cells per cluster (rows) and batch (columns). **d,** UMAP plot of the data with cells coloured by batch. In each batch there is a balanced distribution of cells in the two conditions and across the five clusters. **e,** The Pearson’s gamma (left panel) and the Average Silhouette Width (right panel) was calculated for each set of clusters obtained with 100 random subsamples of 60% of highly variable genes and different values of the deepSplit parameter (see Methods). The most robust clusters correspond to deepSplit values of 0 and 1. **f,** The changes in composition and number of clusters between the clustering obtained with deepSplit 0 (top) and 1 (bottom) are shown using the library “clustree”^57^.

**Extended Data Fig. 2.**
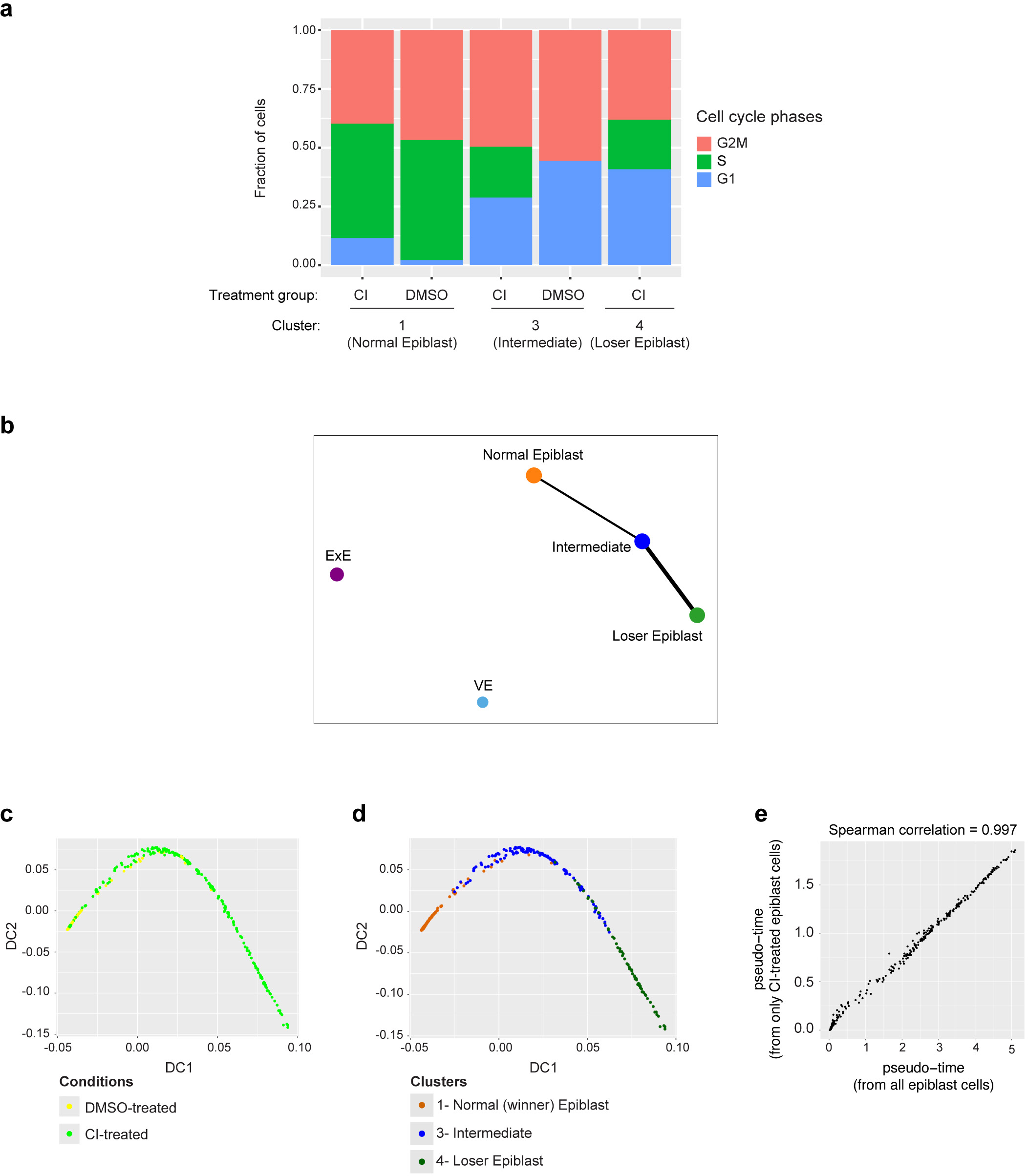
Cell cycle analysis and cluster connectivity. **a,** Cell cycle analysis of epiblast cells from clusters 1, 3 and 4. Cell cycle phase was predicted with cyclone algorithm^58^ and shows that there are cells in S and G2M phase also in the loser and intermediate clusters. **b,** PAGA plot showing the connectivity of the five clusters of cells from CI-treated embryos. **c-d,** Diffusion map analysis in all epiblast cells (from DMSO and CI-treated embryos): cells are coloured according to the condition (**c**) and to the cluster (**d**). **e,** The pseudo-time coordinate of the CI-treated epiblast cells obtained from the diffusion map including all epiblast cells correlates extremely well with the pseudo-time coordinate obtained in the diffusion map calculated only from CI-treated epiblast cells (Fig. 2a).

**Extended Data Fig. 3.**
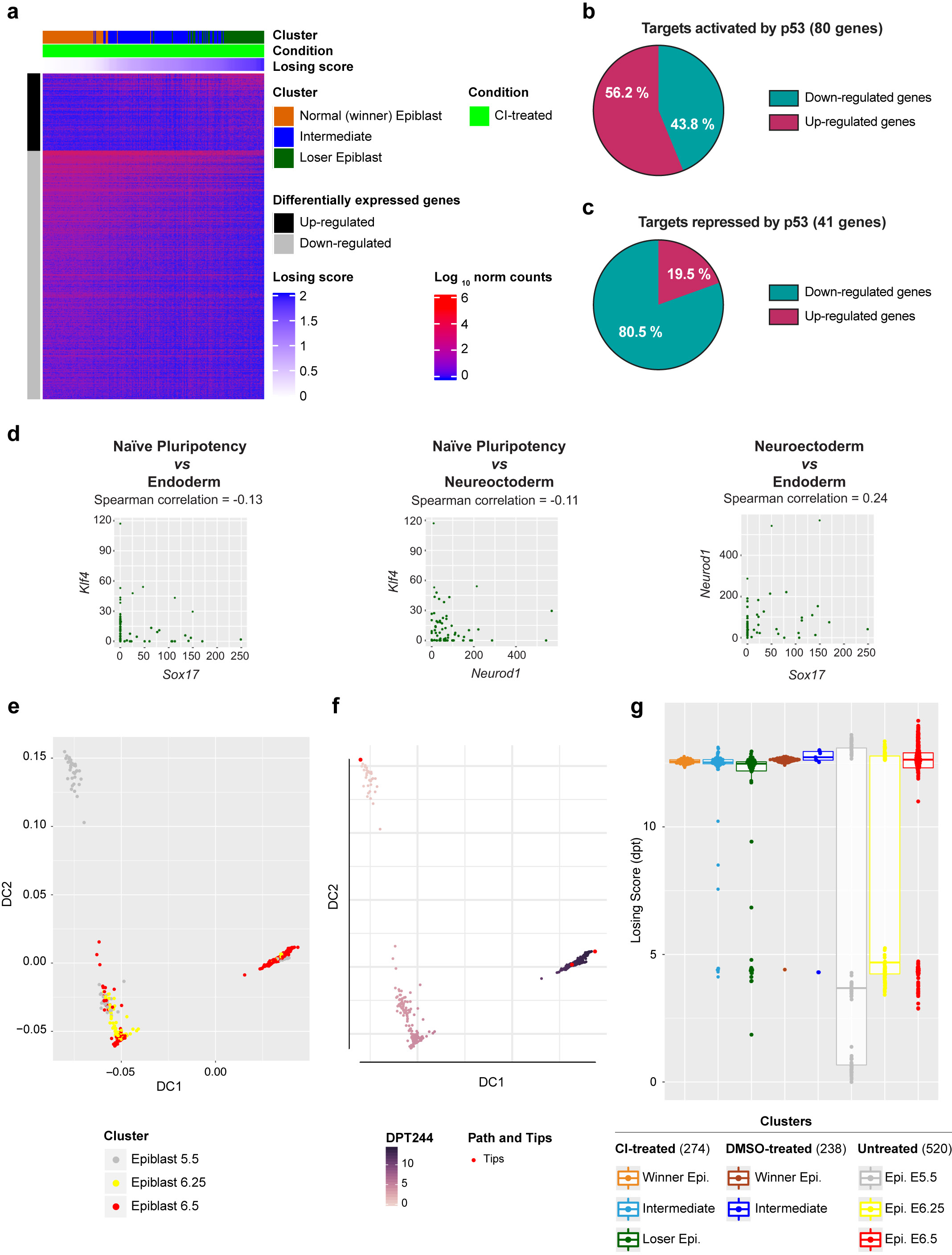
Analysis on epiblast cells from DMSO and CI-treated embryo. **a,** Heatmap showing the expression pattern of all genes differentially expressed along the trajectory from winning to losing cells in Fig. 2d. **b-c,** Overlap of genes differentially expressed along the trajectory joining winning and losing epiblast cells in CI-treated embryos (Fig. 2a and panel d) and genes targeted by p53. Pie charts show the percentage of genes up- or down- regulated in loser cells within the group of target genes that are activated (**b**) or repressed (**c**) by p53. There is an enrichment of activated/repressed targets among genes upregulated/downregulated in losing cells respectively (Fisher’s test, p-value=1E-4). The list of p53 targets is taken from^59^. **d,** Scatter plots of the expression levels of different marker genes plotted against each other in loser epiblast cells (cluster 4). Loser cells have higher expression of pluripotency markers as well as higher expression of some lineage-specific markers and the co-expression of these markers is only weakly correlated. **e-g** Our scRNA-seq data from epiblast cells is projected on top of previously published data from epiblast collected from freshly isolated embryos at different stages (E5.5, E6.25 and E6.5; data from^26^). First, a diffusion map (**e**) and a pseudotime coordinate (**f**) is computed for the epiblast cells from freshly isolated embryos. Then, a pseudotime coordinate is estimated for our data after projecting it onto the diffusion map. Panel **g** shows the pseudotime coordinates for both datasets, split by stage, treatment and cluster.

**Extended Data Fig. 4.**
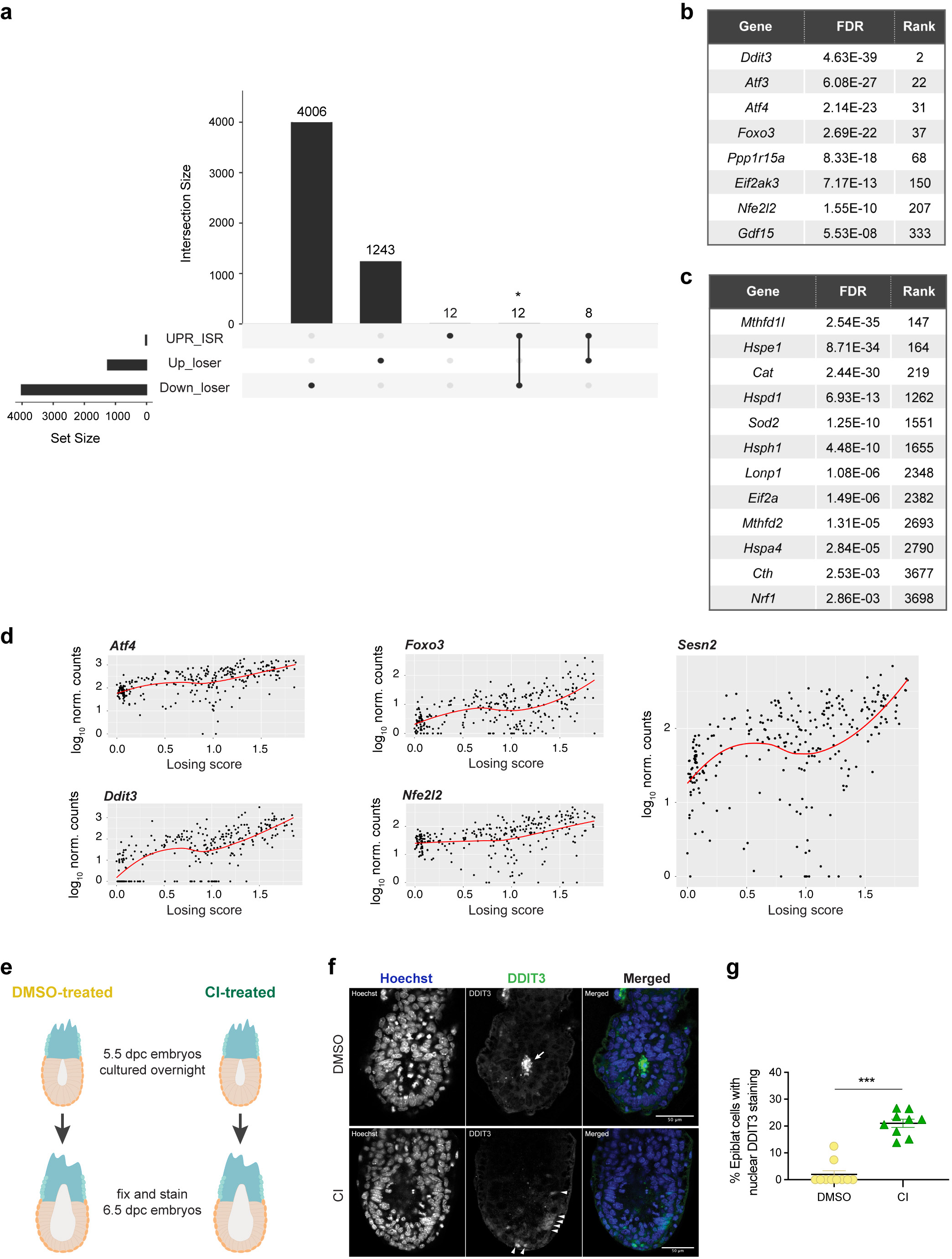
Cells eliminated during early mouse embryogenesis have activated stress responses. **a**, Overlap of genes differentially expressed along the trajectory joining winning and losing epiblast cells in CI-treated embryos (Fig. 2a and Extended Data Fig. 3a) and genes related to the unfolded protein response and integrated protein response pathways (UPR_ISR, see Supplementary Table 3). From the 32 genes related to the UPR & ISR pathways, 12 are down-regulated in loser cells, 8 genes are up-regulated in loser cells, and 12 genes are not differentially expressed between loser and winner cells. There is a statistically significant enrichment of UPR&ISR genes among the up-regulated genes in loser cells (Fisher test, odds ratio=3.0, p-value=0.012). The intersection between UPR-ISR genes and the down regulated genes is not significant (Fisher test, odds ratio=1.2, p value=0.69). **b-c,** List of genes from UPR-ISR pathways that are statistically significantly up-regulated (**b**) or down-regulated (**c**) in loser cells. **d,** Scatterplots with the expression levels of genes involved in stress responses in epiblast cells from CI-treated embryos as a function of cells’ losing score. **e,** Experimental design with the approach taken to validate the expression of the stress response marker DDIT3 in epiblast cells from DMSO or CI-treated embryos. **f,** Representative micrographs of DMSO (upper panel) or CI-treated embryos (100 μM, lower panel) stained for DDIT3, quantified in (**g**). Nuclei are labelled with Hoechst. In control embryos (DMSO-treated), dying cells in the cavity show very high DDIT3 expression (arrow), while live cells in the epiblast of the CI-treated embryos show more modest levels of DDIT3 expression (arrowheads). Scale bar = 20 μm. **g,** Quantification of the percentage of epiblast cells with nuclear DDIT3 expression. N=10 DMSO and N=9 CI-treated embryos. Data shown as mean ± SEM. *Ddit3 (Chop)*, DNA-damage inducible transcript 3; *Atf3*, activating transcription factor 3; *Atf4*, activating transcription factor 4; *Foxo3*, forkhead box O3; *Ppp1r115a (Gadd34),* Protein Phosphatase 1 Regulatory Subunit 15A, *Eif2ak3 (Perk),* Eukaryotic Translation Initiation Factor 2 Alpha Kinase 3; *Nfe2l2 (Nrf2)*, NFE2-related factor 2;. *Sesn2*, Sestrin 2; *Gdf15*, Growth Differentiation Factor 15; *Mthfd1l*, Methylenetetrahydrofolate Dehydrogenase (NADP^+^ Dependent) 1 Like; *Hspe1*, Heat Shock Protein Family E (*Hsp10*) Member 1; *Cat*, Catalase; *Hspd1*, Heat Shock Protein Family D (Hsp60) Member 1; *Sod2,* Superoxide Dismutase 2; *Hsph1,* Heat Shock Protein Family H (*Hsp110*) Member 1; *Lonp1*, Lon Peptidase 1, Mitochondrial; *Eif2a,* Eukaryotic Translation Initiation Factor 2A; *Mthfd2*, Methylenetetrahydrofolate Dehydrogenase (NADP^+^ Dependent) 2, Methenyltetrahydrofolate Cyclohydrolase; *Hspa4*, Heat Shock Protein Family A (*Hsp70*) Member 4; *Cth*, Cystathionine Gamma-Lyase; *Nrf1*, Nuclear Factor 1.

**Extended Data Fig. 5.**
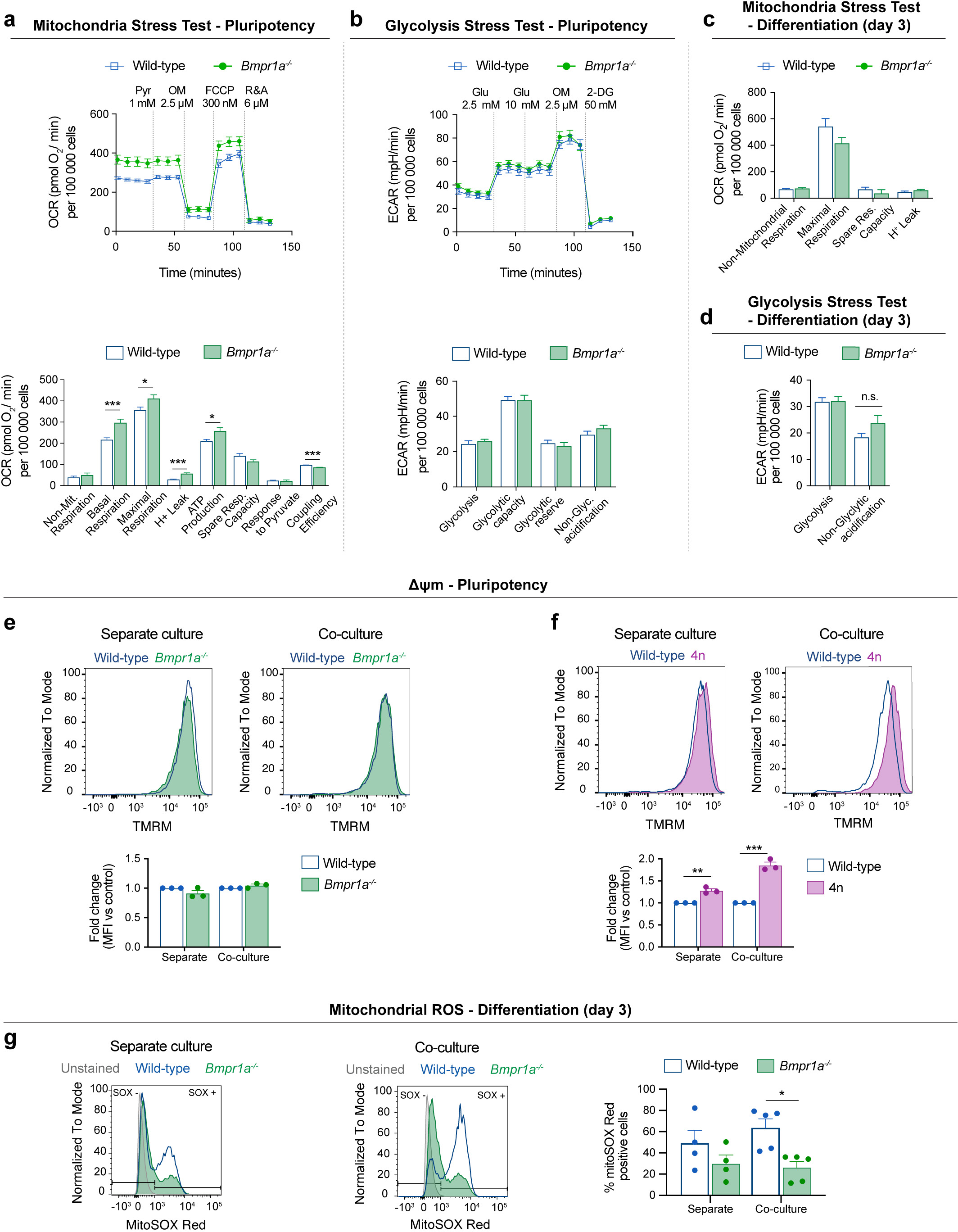
Mitochondrial function in Wild-type, *Bmpr1a^-/-^* and 4n mESCs. **a-d,** Metabolic flux analysis of wild-type and *Bmpr1a^-/-^* mESCs. OCR profile and metabolic parameters assessed during the mitochondria stress test performed in pluripotency conditions (**a**). ECAR profile and metabolic parameters assessed during the glycolysis stress test performed in pluripotency conditions (**b**). Metabolic parameters from the mitochondria stress test found to be similar between wild-type and *Bmpr1a^-/-^* mESCs during differentiation – day 3 (**c**). Metabolic parameters from the glycolysis stress test found to be similar between wild-type and *Bmpr1a^-/-^* mESCs during differentiation – day 3 (**d**). Data obtained with a minimum of 3 independent experiments, with 5 replicates per cell type in each assay. Statistical analysis done with Mann-Whitney test. **e-f,** Analysis of mitochondrial membrane potential (Δψm) in defective mESCs maintained in pluripotency conditions, in separate or co-culture. Representative histograms of TMRM fluorescence and quantification for wild-type and *Bmpr1a^-/-^* (**e**) and wild-type and 4n (**f**). Statistical analysis done by two-way ANOVA, followed by Holm-Sidak’s multiple comparisons test. **g,** Analysis of mitochondrial ROS in wild-type and *Bmpr1a^-/-^* mESCs undergoing differentiation in separate or co-culture: representative histograms of mitoSOX Red fluorescence and quantification of the percentage of mitoSOX positive cells. Statistical analysis done by two-way ANOVA, followed by Holm-Sidak’s multiple comparisons test. Data obtained with a minimum of 3 independent experiments. Error bars represent SEM.

**Extended Data Fig. 6.**
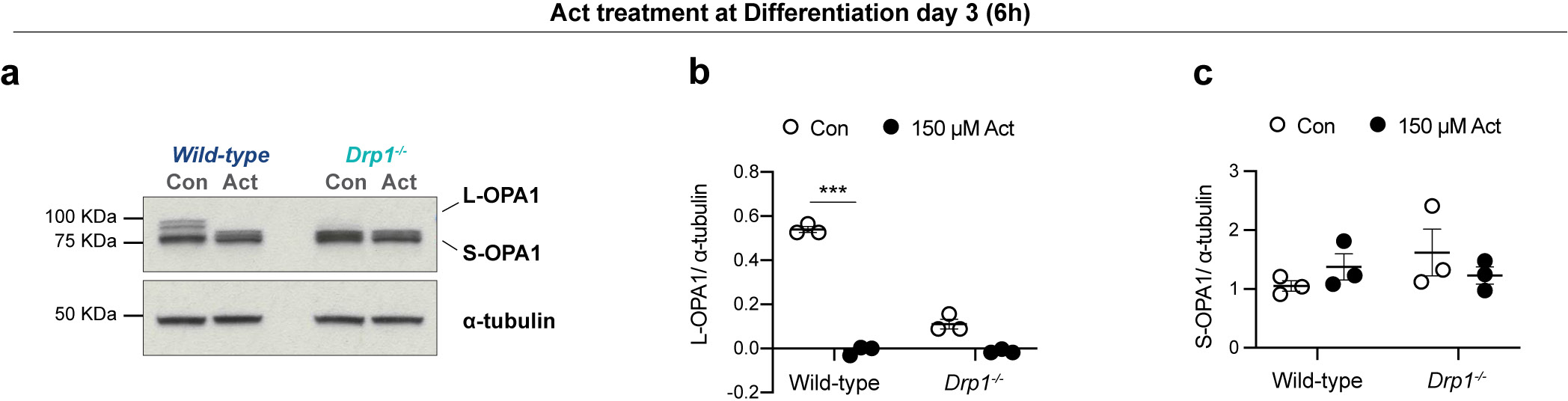
Effect of actinonin in OPA1 expression in wild-type and *Drp1^-/-^* cells. **a**, Western blot analysis of OPA1 expression in wild-type and *Drp1^-/-^* cells treated with actinonin (Act, 150 μM) during 6 hours on the third day of differentiation, quantified in (**b-c**). **b-c**, Expression levels of L-OPA1 (**b**) and S-OPA1 (**c**) relative to ɑ-tubulin. Data shown as mean ± SEM of a minimum of 3 independent experiments. Statistical analysis done by two-way ANOVA, followed by Holm-Sidak’s multiple comparisons test.

**Extended Data Fig. 7.**
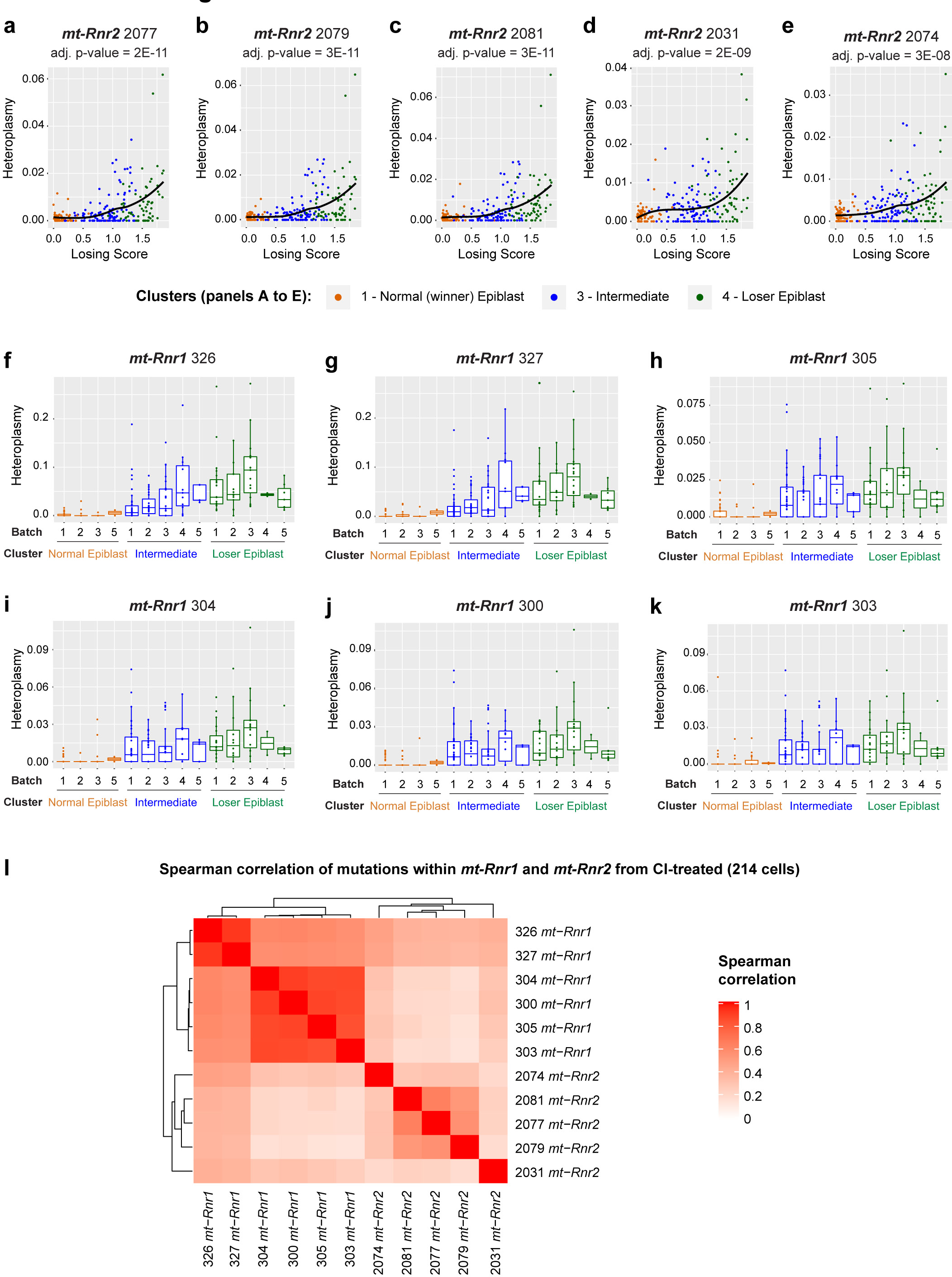
Analysis of SNPs in mtDNA in epiblast cells. **a-e,** mtDNA heteroplasmy in epiblast cells from CI-treated embryos for five positions within the *mt-Rnr2* gene. All of these positions have an heteroplasmy that increases with the cells’ losing scores in a statistically significant way (the adjusted p-value estimated via a generalized linear model is indicated at the top of each plot). **f-k,** The variation in the heteroplasmy across the CI-treated cells is not due to a batch effect for the 6 significant positions within the *mt-Rnr1* gene. **l,** Spearman’s correlation between the mtDNA heteroplasmy at all the statistically significant positions (six within the gene mt-*Rnr1* and five within the gene mt-*Rnr2*).

**Extended Data Fig. 8.**
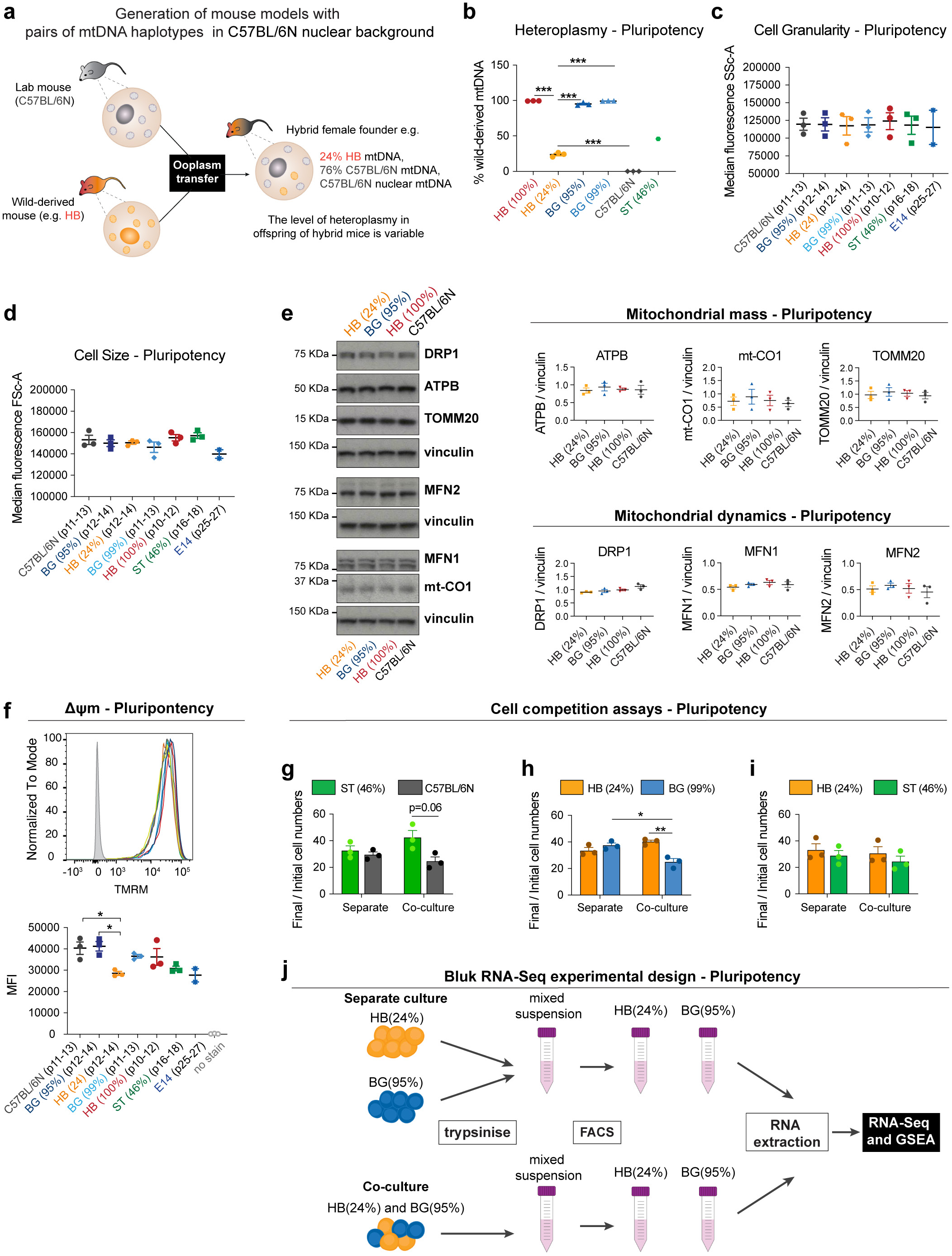
Changes in mtDNA sequence are enough to trigger cell competition. **a**, Illustration of the process of derivation of the mESCs lines from mice that are hybrid between the wild-caught strains (BG, HB or ST) and the lab mouse (C57BL/6N). These hybrid mice were generated elsewhere^16^ by ooplasmic transfer: the zygote of a C57BL/6N mouse was injected with ooplasm from a wild-caught mouse (orange, HB pictured). Therefore, these hybrid mice contain the nuclear background of the C57BL/6N strain and the mtDNA of wild-caught strain and potentially C57BL/6N mtDNA (heteroplasmic mice strains). mESCs lines were derived from the hybrid mice and characterised. **b-f,** Characterisation of the derived cell lines by flow cytometry, during pluripotency, in comparison to the wild-type cell line used in previous experiments (E14, 129/Ola background). Heteroplasmy analysis of the derived mESC lines from the hybrid mice, indicating the percentage of wild-derived mtDNA (**b**). Cell granularity (internal complexity) given as median fluorescence intensity of SSc-A laser (**c**). Cell size given as median fluorescence intensity of FSc-A laser (**d**). Analysis of the expression of mitochondrial markers: representative western blot and quantification of markers of mitochondrial mass (ATPB, mt-CO1 and TOMM20) and mitochondrial dynamics (DRP1, MFN1and MFN2), relative to vinculin, in cells derived from hybrid mice (**e**). **f,** Representative histograms and quantification of median TMRM fluorescence, indicative of Δψm, for the hybrid cell lines derived, in comparison to the wild-type cell line used in previous experiments (E14, 129/Ola background). Statistical analysis done by one-way ANOVA, followed by Holm-Sidak’s multiple comparisons test. **g-i,** Cell competition assays between hybrid cell lines maintained in pluripotency culture conditions. The ratio of final/initial cell numbers in separate or co-culture is shown. Statistical analysis done by two-way ANOVA, followed by Holm-Sidak’s multiple comparisons test. **j,** Experimental design for RNA-Seq and gene set enrichment analysis (GSEA). The isolation of RNA from winner HB(24%) and loser BG(95%) cells was performed after three days in separate or co-culture conditions, once cells have been subjected to FACS to isolate the two populations form mixed cultures. Data obtained with a minimum of 3 independent experiments. Error bars represent SEM.

**Extended Data Fig. 9.**
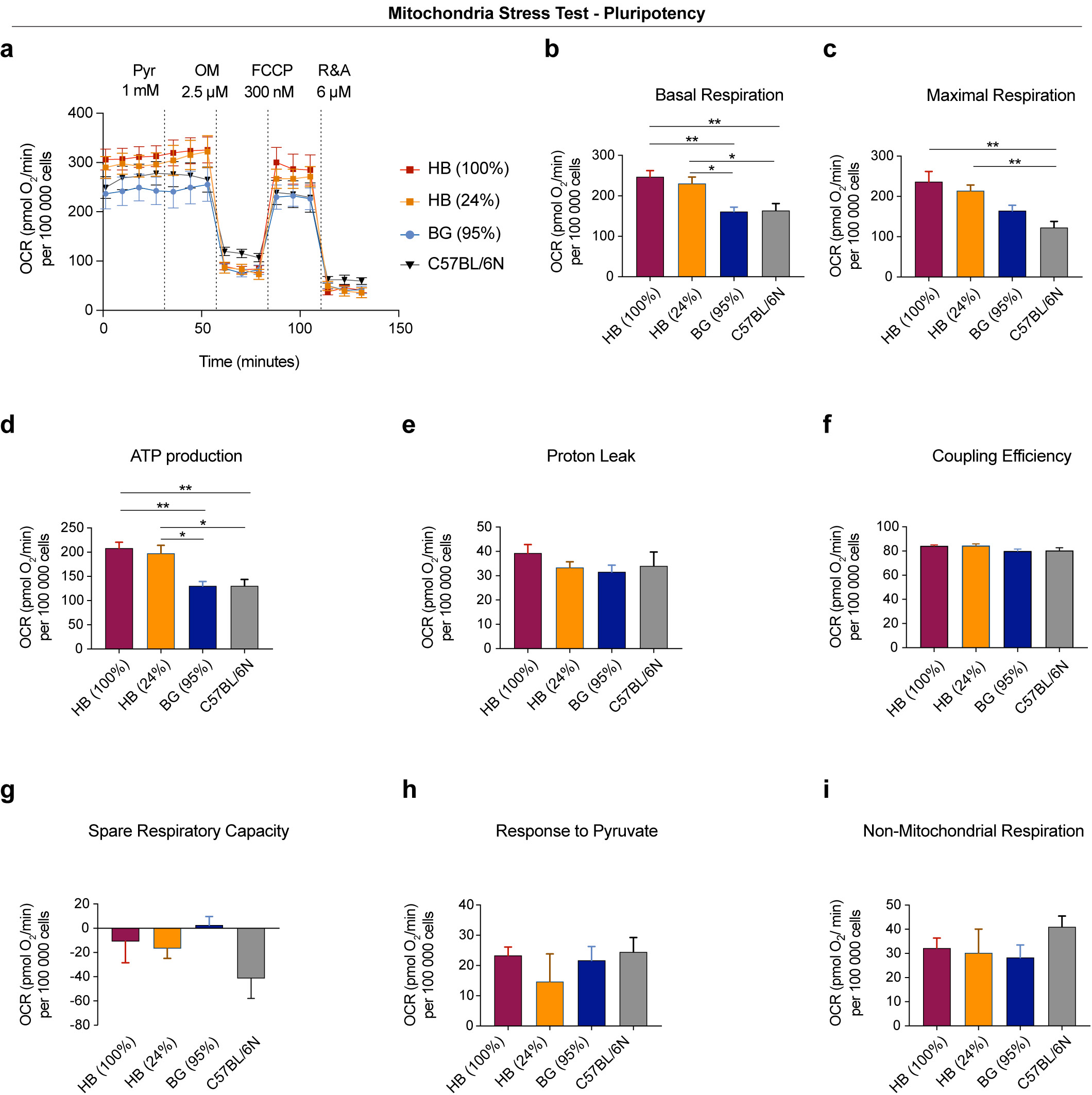
Metabolic flux analysis of the cells with different mtDNA variants: HB(100%), HB(24%), BG(95%) and C57BL/6N. **a,** OCR profile during mitochondria stress test performed in pluripotency maintenance conditions. **b-i,** Metabolic parameters assessed during the during the mitochondria stress test performed in pluripotency conditions. Data obtained with a minimum of 3 independent experiments, with 5 replicates per cell type in each assay. Error bars represent SEM. Statistical analysis done with Kruskal-Wallis test, followed by Dunn’s multiple comparison test.

**Extended Data Fig. 10.**
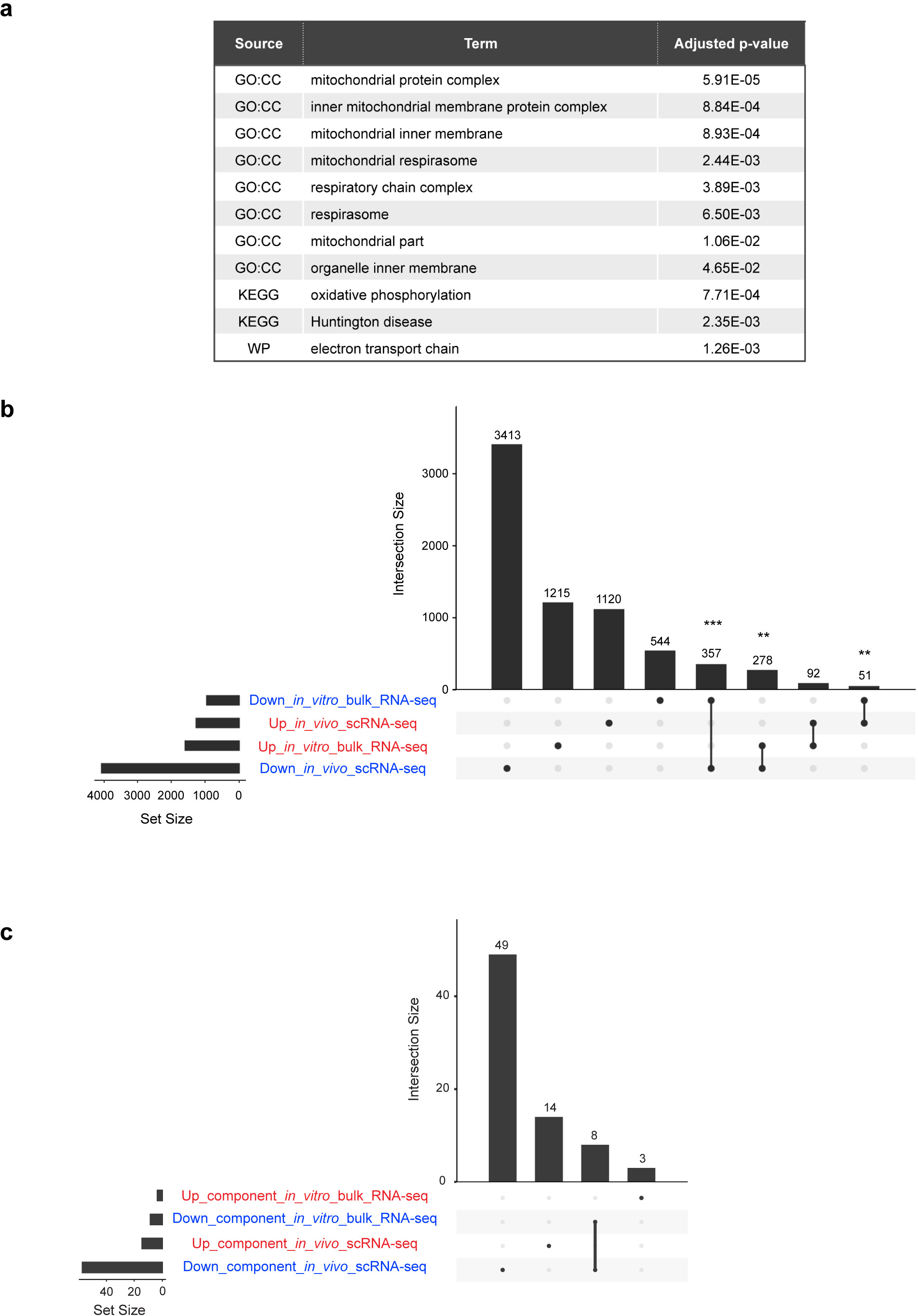
Common features of scRNA-seq and bulk RNA-seq datasets. **a,** Terms significantly enriched among genes downregulated in BG(95%) (loser) ESCs *in vitro* when co-cultured with HB(24%) cells. The loss of mitochondrial activity emerges as a common feature between loser cells *in vivo* and *in vitro*. The gene enrichment analysis was performed using g-profiler tool (see Methods). **b,** Intersection between differentially expressed genes along the trajectory from winning to losing epiblast cells (“in_vivo_scRNA-seq”; Fig. 2a and Extended Data Fig. 3a and genes differentially expressed between co-cultured HB(24%) (winner) and BG(95%) (loser) ESCs (“in_vitro_bulk_RNA-seq”). “Up” and “Down” here refer to genes up- or down-regulated in loser cells. Fisher test for the intersection between down-regulated genes from scRNA-seq (*in vivo*) and down-regulated genes from bulk RNA-seq (*in vitro*): p-value, 1.71E-12; odds ratio 1.80. Fisher test for the intersection between down-regulated genes from scRNA-seq (*in vivo*) and up-regulated genes from bulk RNA-seq (*in vitro*): p-value, 5.20E-3; odds ratio 0.67. Fisher test for the intersection between up-regulated genes from scRNA-seq (*in vivo*) and down-regulated genes from bulk RNA-seq (*in vitro*): Fisher test p-value, 4.87E-3; odds ratio 0.80. The intersection between up-regulated genes from sc-RNA-seq (*in vivo*) and up-regulated genes from bulk RNA-Seq (*in vitro*) is not statistically significant: Fisher test p-value: 0.30, odds ratio 1.14. **c,** Intersection between the significantly enriched terms in genes upregulated or downregulated in loser cells in the epiblast of CI-treated embryos (“*in_vivo*_scRNA-Seq”) or in our *in vitro* model of competition between co-cultured HB(24%) (winner) and BG(95%) (loser) ESCs (“*in_vitro*_bulk_RNA-seq”). All the terms enriched among downregulated genes *in vitro* are also enriched *in vivo*.

## List of Tables

**Supplementary Table 1**. List of genes down-regulated along the winner-to-loser trajectory in the embryo.

**Supplementary Table 2**. List of genes up-regulated along the winner-to-loser trajectory in the embryo.

**Supplementary Table 3**. Genes related to the unfolded protein response and integrated protein response pathways (UPR_ISR) that were analysed in the genes differentially expressed along the winner-to-loser trajectory.

**Supplementary Table 4**. List of background genes for the winner-to-loser trajectory in the embryo.

**Supplementary Table 5**. List of genes down-regulated in BG(95%) cells when co-cultured with HB(24%) cells.

**Supplementary Table 6**. List of genes up-regulated in BG(95%) cells when co-cultured with HB(24%) cells.

**Supplementary Table 7**. List of background genes used for the analysis of genes differentially expressed between co-cultured BG(95%) and HB(24%) cells.

## Methods

### Animals

Mice were maintained and treated in accordance with the Home Office’s Animals (Scientific Procedures) Act 1986 and covered by the Home Office project license PBBEBDCDA. All mice were housed on a 10 hr-14 hr light-dark cycle with access to water and food *ad libitum*. Mattings were generally set up in the afternoon. Noon of the day of finding a vaginal plug was designated embryonic day 0.5 (E0.5). Embryo dissection was performed at appropriate timepoints in M2 media (Sigma), using Dumont No.5 forceps (11251-10, FST). No distinction was made between male and female embryos during the analysis.

### Cell lines, cell culture routine and drug treatments

E14, kindly provided by Prof A. Smith, from Cambridge University, were used as wild-type control cells tdTomato-labelled or unlabelled. GFP-labelled or unlabelled cells defective for BMP signalling (*Bmpr1a^-/-^*), tetraploid cells (4n) and *Bmp1a^-/-^* null for p53 *(Bmpr1a^-/-^;p53*^-/*-*^) are described elsewhere ^6, 7^. Cells null for Dynamin-related protein 1 (*Drp1^-/-^*) or Mitofusin 2 (*Mfn2^-/-^*) were generated by CRISPR mutagenesis. Cells with different mitochondrial DNA (mtDNA) content in the same nuclear background were derived from embryos of hybrid mice, generated elsewhere ^16^.

Cells were maintained pluripotent and cultured at 37°C in 5% CO_2_ in 25 cm^2^ flasks (Nunc) coated with 0.1% gelatin (Sigma) in DPBS. Growth media (ES media) consisted of GMEM supplemented with 10% FCS, 1 mM sodium pyruvate, 2 mM L-glutamine, 1X minimum essential media non-essential amino-acids, 0.1 mM β-mercaptoethanol (all from Gibco) and 0.1% leukemia inhibitory factor (LIF, produced and tested in the lab). Cells derived from hybrid mice (C57BL/6N nuclear background) were maintained on 0.2% LIF. The growth media was changed daily, and cells were split every 3 days.

To manipulate mitochondrial translation during differentiation, wild-type and *Drp1^-/-^* mESCs were treated with doxycycline (Dox, 22.5 μM), from day 1 to day 3 of culture, or with actinonin (Act, 150 μM), for 6 hours on day 3 of culture in N2B27 media (see Differentiation and Cell competition assays). As control condition, cells were treated with vehicle (Con). Samples were collected on day 3 of differentiation for western blot analysis.

### CRISPR mutagenesis

Drp1 and Mfn2 knockout ESCs were generated by CRISPR-Cas9 mediated deletion of Drp1 exon 2 and Mfn2 exon 3 respectively. sgRNA guides flanking Drp1 exon 2 or Mfn2 exon 3 were cloned into the PX459 vector (Addgene)^60^: Drp1 exon 2 upstream sgRNA: 5’ TGGAACGGTCACAGCTGCAC 3’; Drp1 exon 2 downstream sgRNA: 5’ TGGTCGCTGAGTTTGAGGCC 3’; Mfn2 upstream sgRNA: 5’ GTGGTATGACCAATCCCAGA 3’; Mfn2 downstream sgRNA: 5’ GGCCGGCCACTCTGCACCTT 3’. E14 ESCs were co-transfected with 1ug of each sgRNA expression using Lipofectamine 2000 (Invitrogen) according to manufacturer’s instructions. As control E14 ESCs were transfected in parallel with equal amount of empty PX459 plasmid. Following 6 days of Puromycin selection, single colonies were picked from both Drp1 sgRNA and empty vector transfected ESCs and screened for mutations. Drp1 exon 2 deletion was confirmed by PCR genotyping using the following primers: Drp1_genot F: 5’ GGATACCCCAAGATTTCTGGA 3’; Drp1_genot R: 5’ AGTCAGGTAATCGGGAGGAAA 3’, followed by Sanger Sequencing. Mfn2 exon 3 deletion was confirmed by PCR genotyping using the following primers: Mfn2_genot F: 5’ CAGCCCAGACATTGTTGCTTA 3’; Mfn2_genot R: 5’ AGCTGCCTCTCAGGAAATGAG 3’, followed by Sanger Sequencing.

### Derivation of mESCs from hybrid mouse strains and heteroplasmy determination

The derivation of new mESC lines was adapted from ^61^. Cells were derived from embryos of hybrid mouse strains BG, HB and ST. These contain the mtDNA of C57BL/6N (Bl6) lab mouse and mtDNA variants from wild-caught mice ^16^.

Embryos were isolated at E2.5 (morula stage) and cultured in 4-well plates (Nunc, Thermo Scientific) containing KSOM media (Millipore) plus two inhibitors (KSOM+2i): 1 μM MEK inhibitor PDO325901 (Sigma-Aldrich) and 3 μM GSK-3 inhibitor CHIR9902 (Cayman Chemicals) for 2 days at 37°C in 5% CO_2_ incubator. To reduce evaporation, the area surrounding the wells was filled with DPBS. Embryos were further cultured in a fresh 4-well plates containing.

N2B27+2i+LIF media: N2B27 media supplemented with 1 μM MEK inhibitor PDO325901 and 3 μM GSK-3 inhibitor and 0.1% LIF for up to 3 days until reaching the blastocyst stage. Each embryo was then transferred to a well of a 96-well plate coated with 0.1% gelatin in DPBS and containing 150 μL of N2B27+2i+LIF media per well. In these conditions, the embryos should attach to the wells allowing the epiblast to form an outgrowth. This plate was then incubated at 37°C in 5% CO_2_ incubator for 3 to 7 days until ES-like colonies start to develop from the epiblast outgrowth. Cells were passaged by dissociation with Accutase (Sigma) and seeded in gradual increasing surface area of growth (48-well, 24-well, 12-well plate, T12.5 and T25 flask), until new cell lines were established. At this stage cells were weaned from N2B27+2i+LIF media and then routinely cultured in ES media.

These new cell lines were then subjected to characterisation by flow cytometry (cell size, granularity and mitochondrial membrane potential) and ARMS-qPCR assay^16^ (to determine heteroplasmy).

### Embryo experiments

Early mouse embryos were isolated at E5.5 (from pregnant CD1 females, purchased from Charles River, UK). Following dissection from the decidua, embryos were cultured overnight in N2B27 “poor” media (same formulation as N2B27 media but supplemented with 0.5xB27 supplement and 0.5xN2 supplement) with pan-caspase inhibitors (100 μM, Z-VAD-FMK, FMK001, R&D Systems, USA) or equal volume of vehicle (DMSO) as control. On the next morning, embryos were processed for single cell RNA-Seq (scRNA-seq) or functional validation (Δψm analysis and immunohistochemistry for markers of loser cells).

For the scRNA-seq and Δψm analysis embryos were dissociated into singe-cells. Briefly, up to 12 embryos were dissociated in 600 μL Acccutase (A6964, Sigma, UK) during 12 min at 37°C, tapping the tube every two minutes. Accutase was then neutralised with equal volume of FCS, cells span down and stained with TMRM, for Δψm analysis, or directly re-suspended in 300 μL DPBS with 1% FCS, for single cell sorting and RNA-seq. Sytox blue (1:1000, S34857, ThermoFisher Scientific, UK), was used as viability staining.

### Differentiation and Cell competition assays

Cell competition assays between wild-type and *Bmpr1a^-/-^*, 4n or *Drp1^-/-^* cells were performed in differentiating conditions. Cells were seeded onto fibronectin-coated plates (1:100, Merck) in DPBS during 1h at 37°C and grown in N2B27 media - to promote the differentiation of mESCs into a stage resembling the post-implantation epiblast, as cell competition was previously shown to occur in these conditions ^6^. N2B27 media consisted of 1:1 Dulbecco’s modified eagle medium nutrient mixture (DMEM/F12) and Neurobasal supplemented with N2 (1x) and B27 (1x) supplements, 2 mM L-glutamine and 0.1 mM β-mercaptoethanol - all from Gibco. Cell competition assays between wild-type and *Mfn2^-/-^* and between mESCs with different mtDNA content were performed in pluripotency maintenance conditions (ES media).

Cells were seeded either separately or mixed for co-cultures at a 50:50 ratio, onto 12 well plates, at a density of 8E04 cells per well, except for assays between wild-type and *Mfn2^-/-^* mESCs, where 3.2E05 cells were seeded per well. The growth of cells was followed daily and compared between separate or co-culture, to control for cell intrinsic growth differences, until the fourth day of culture. Viable cells were counted daily using Vi-CELL XR Analyser (Beckman Coulter, USA), and proportions of each cell type in co-cultures were determined using LSR II Flow Cytometer (BD Bioscience), based on the fluorescent tag of the ubiquitously expressed GFP or TdTomato in one of the cell populations.

### Metabolomic analysis

The metabolic profile was obtained using the Metabolon Platform (Metabolon, Inc). Each sample consisted of 5 biological replicates. For each replicate, 1E07 cells were spun down and snap frozen in liquid nitrogen. Pellets from 5 independent experiments for each condition were analysed by Metabolon Inc by a combination of Ultrahigh Performance Liquid Chromatography-Tandem Mass Spectroscopy (UPLC-MS/MS) and Gas Chromatography-Mass Spectroscopy (GC-MS). Compounds were identified by comparison to library entries of purified standards based on the retention time/index (RI), mass to charge ratio (*m/z)*, and chromatographic data (including MS/MS spectral data) on all molecules present in the library. Samples were normalized to protein content measured by Bradford assay. Statistical analysis was done using Welch’s two-sample t-test and statistical significance defined as p ≤0.05.

### Seahorse analysis

The metabolic function of cells was assessed by extracellular flux analysis using Seahorse XF24 (Agilent Technologies, UK). For assays ran during pluripotency, cells were seeded, on the day prior to the assay, onto 0.1% gelatin-coated (Sigma, UK) in 300 µL of ES media. All cell types were seeded at 5×10^4^ cells per well, except for *Bmpr1a^-/-^* cells, that were seeded at 6E04 per well). For assays ran during differentiation, cells were seeded, the 3 days before the assay, onto fibronectin-coated fibronectin-coated plates (1:100, Merck, UK), in 300 µL of N2B27media. All cell types were seeded at 2.4E04 cells per well, except for *Bmpr1a^-/-^* cells, that were seeded at 3.2E04 cells per well.

On the day of the assay, cells were carefully washed twice with assay media and then left with a final volume of the 600 µL per well. The plate was then equilibrated on a non-CO_2_ incubator at 37°C for 30 min. The assay media consisted in unbuffered DMEM (D5030 – Sigma, UK), that was supplemented on the day of the assay according to the test performed. For the OCR measurements the assay media was supplemented with 0.5 g.L−1 of glucose (Sigma, UK) and 2 mM of L-glutamine (Life Technologies, UK), while for the ECAR measurements the media was supplemented with 1 mM of Sodium Pyruvate and 2 mM of L-glutamine (both from Life Technologies, UK), pH 7.4 at 37°C.

The protocol for the assay consisted of 4 baseline measurements and 3 measurements after each compound addition. Compounds (all from Sigma, UK) used in OCR and ECAR assays were prepared in the supplemented assay media. For the OCR assay, test the following compounds were added: 1 mM Pyruvate (Pyr), 2.5 µM oligomycin (OM), 300 nM Carbonyl cyanide-4-(trifluoromethoxy) phenylhydrazone (FCCP) and a mixture of rotenone and antimycin A at 6 µM each (R&A). For the ECAR assay, the following compounds were added: 2.5 mM and 10 mM of glucose, 2.5 µM of oligomycin (OM), and a 50 mM of 2-deoxyglucose (2-DG).

Each of the experiments was performed in 3 times, with 5 biological replicates of each cell type. For background correction measurements, 4 wells were left without cells (A1, B4, C3 and D6). Both ECAR and OCR measurements were performed on the same plate. The assay parameters for both tests were calculated following the Seahorse assay report generator (Agilent Technologies, UK).

At the end of the assay, cells were fixed and stained with Hoechst. Both OCR and ECAR were normalised to cell number, determined by manual cell counts using Fiji software. The normalisation of the data was processed on Wave Desktop software (Agilent Technologies, UK) and data exported to Prism 8 (GraphPad) for statistical analysis.

### Analysis of mitochondrial membrane potential (Δψm) and mitochondrial ROS

Quantitative analysis of Δψm and mitochondrial ROS was performed by flow cytometry. Cells were grown in pluripotency or differentiating conditions, as described above. Cells were dissociated and pelleted to obtain 2E05 cells per sample for the staining procedure.

For TMRM staining in single cells from early mouse epiblasts, embryos were dissected at E5.5 and cultured overnight in the presence or absence of caspase inhibitors. On the following morning, to avoid misleading readings, epiblasts were isolated initially by an enzymatic treatment with of 2.5% pancreatin, 0.5% trypsin and 0.5% polyvinylpyrrolidone (PVP40) - all from Sigma-Aldrich- to remove the visceral endoderm (VE). Embryos were treated during 8 min at 4°C, followed by 2 min at RT. The VE was then pealed with the forceps and the extraembryonic ectoderm removed to isolate the epiblasts. Up to 16 epiblasts were pooled per 600µL of Accutase (Sigma-Aldrich) for dissociation into single cells prior to staining. Reaction was stopped with equal volume of FCS and cells subjected to TMRM staining.

Cells were loaded with 10 nM of the Nernstian probe tetramethylrhodamin methyl ester perchlorate (TMRM, Sigma), prepared in N2B27 media. After incubating for 15 min at 37°C, cells were pelleted again and re-suspended in flow cytometry (FC) buffer (3% FCS in DPBS). Sytox blue (1:1000, Invitrogen, UK) was used as viability staining. Stained cell suspensions were analysed in BD LSRII flow cytometer operated through FACSDiva software (Becton Dickinson Biosciences, UK). For TMRM fluorescence detection the yellow laser was adjusted for excitation at λ=562 nm, capturing the emission light at λ=585 nm for TMRM. In the case of GFP-labelled cell lines, for GFP fluorescence detection the blue laser was adjusted for excitation at λ=488 nm, capturing the emission light at λ=525 nm. Results were analysed in FlowJo vX10.0.7r2.

Qualitative analysis of Δψm was performed by confocal microscopy. Wild-type and *Bmpr1a^-/-^* cells were grown in fibronectin-coated glass coverslips. On the third day of differentiation, cells were loaded with 200 nM MitoTracker Red probe (Life Technologies), prepared in N2B27 media, for 15 min at 37°C. Cells were then washed with DPBS and fixed with 3.7% formaldehyde for subsequent immunocytochemical staining of total mitochondria mass, with TOMM20 antibody.

For the analysis of mitochondrial ROS, cells were grown in differentiating conditions and stained on the third day of culture. Briefly, 2E05 cells of each cell line were resuspended in 200 μL of 5 μM solution of MitoSOX (Invitrogen, UK) prepared in N2B27 media. Cells were incubated at 37°C for 15 min, and then resuspended in FC buffer. MitoSOX fluorescence was analysed with the violet laser adjusted for excitation at λ=405 nm, capturing the emission light at λ=610 nm. Sytox blue was used as viability staining.

### Immunofluorescence

Cells were washed with DPBS and fixed with 3.7% formaldehyde (Sigma, UK) in N2B27, for 15 min at 37°C. Permeabilization of the cell membranes was done with 0.4% Triton X-100 in DPBS (DPBS-Tx), at RT with agitation. Blocking step with 5% BSA in DPBS-Tx 0.1% was performed for 30 min, at RT with agitation. Mitochondria were labelled with TOMM20 antibody (1:100,

Santa Cruz Biotechnologies). Dead cells were labelled with cleaved caspase-3 antibody (1:400, CST) and NANOG antibody was used to mark pluripotent cells (1:100, eBioscience). Secondary antibodies were Alexa Fluor (1:600, Invitrogen). Primary antibody incubation was performed overnight at 4°C and secondary antibody incubation during 45 min, together with Hoechst to stain nuclei (1:1000, ThermoScientific), at RT and protected from light. In both cases antibodies were diluted in blocking solution. Three 10 min washes with DPBS-Tx 0.1% were performed between each critical step and before mounting with Vectashield medium (Vector Laboratories).

Samples were imaged with a Zeiss LSM780 confocal microscope (Zeiss, UK) and processed with Fiji software ^62^. Mitochondria stainings were imaged with a 63x/1.4 Oil objective. For samples stained with TOMM20 antibody and MitoTracker Red, Z-stacks were acquired and processed for deconvolution using Huygens software (Scientific Volume Imaging, https://svi.nl/). Samples stained with cleaved caspase-3 were imaged with 20x/0.8 air objective. Imaging and deconvolution analysis were performed with the support and advice from Mr. Stephen Rothery from the Facility for Imaging by Light Microscopy (FILM) at Imperial College London.

Embryo immunofluorescent staining for p-rpS6, OPA1 and DDIT3 (CHOP) markers was performed as follows. Cultured embryos were fixed in 4% PFA in DPBS containing 0.01% Triton and 0.1% Tween 20 during 20 min at RT. Permeabilization of the membranes was done during 10 min in DPBS with 0.5% Triton. Embryos were blocked in 5% BSA in DPBS with 0.25% Triton during 45 min. Incubation with primary antibodies - CHOP (1:500, CST-2895S), OPA1 (1:100, BD Biosciences - 612606) and p-rpS6 (CST - 5364) - was done overnight at 4°C in 2.5% BSA in DPBS with 0.125% Triton. On the following morning, hybridisation with secondary antibodies Alexa Fluor 568 and Alexa Fluor 488 (diluted 1:600 in DPBS with 2.5% BSA and 0.125% Triton) was done next during 1h at RT. Hoechst was also added to this mixture to stain nuclei (1:1000, Invitrogen). Three 10 min washes with filtered DPBS-Tx 0.1% were performed between each critical step. All steps were done with gentle agitation.

Embryos were imaged in embryo dishes (Nunc) in a drop of Vectashield using Zeiss LSM780 confocal microscope at 40x/1.3 oil objective.

Further details about image acquisition and processing are specified in the Supplementary Methods file “Imaging equipment and settings.docx”

### Western Blotting

Cells were washed in DPBS and lysed with Laemmli lysis buffer (0.05 M Tris-HCl at pH 6.8, 1% SDS, 10% glycerol, 0.1% β-mercaptoethanol in distilled water). Total protein quantification was done using BCA assay (Thermo Scientific, UK) and samples (15μg of protein per lane) were loaded into 12% Bis-Tris protein gels (BioRad). Resolved proteins were transferred into nitrocellulose membranes (GE Healthcare). The following primary antibodies were incubated overnight at 4°C: rabbit anti-TOMM20 (1:1000, CST - 42406), rabbit anti-α-Tubulin (1:1000, CST- 2144), mouse anti-mt-CO1 (1:2000, Abcam - 14705), rabbit anti-DRP1 (1:1000, CST-8570), mouse anti-MFN1 (1:1000, Abcam - 57602), mouse anti-MFN2 (1:500, Abcam - 56889), mouse anti-Vinculin (1:1000, Sigma - V9131), mouse anti-OPA1 (1:1000, BD Biosciences - 612606), rabbit anti-ATF4 (1:1000, CST-11815), rabbit anti-α-PCNA (1:1000, Abcam - 2426) and rabbit anti-p-eIF2ɑ (Ser51, 1:1000, CST-9721). On the following morning, HRP-conjugated secondary antibodies (Santa Cruz) were incubated for 1h at RT. Membranes were developed with ECL reagents (Promega) and mounted in cassette for time-time-controlled exposure to film (GE Healthcare).

### Bulk RNA-Seq and Single cell RNA-Seq

For bulk RNA Seq in the competitive scenario between cells with different mtDNA, HB(24%) and BG(95%) mESCs were grown separately or in co-culture. On the third day of culture cells were dissociated and subjected to fluorescence activated cell sorting (FACS) to separate the cell populations in co-culture. To control for eventual transcriptional changes due to the FACS process, a mixture of the two separate populations was subjected to the same procedure as the co-cultured samples. Total RNA isolation was then carried out using RNA extraction Kit (RNeasy Mini Kit, Qiagen). PolyA selection/enrichment was the method adopted for library preparation, using the NEB Ultra II RNA Prep Kit. Single end 50bp libraries were sequenced on Illumina Hiseq 2500. Raw basecall files were converted to fastq files using Illumina’s bcl2fastq (version 2.1.7). Reads were aligned to the mouse genome (mm9) using Tophat2 version 2.0.11 ^63^ with default parameters. Mapped reads that fell on genes were counted using featureCounts from Rsubread package ^64^. Generated count data were then used to identify differentially expressed genes using DESeq2 ^65^. Genes with very low read counts were excluded. Finally, Gene Set Enrichment Analysis was performed using GSEA software ^66, 67^ on pre-ranked list generated by DESeq2.

To investigate the nature of cells eliminated by cell competition during early mouse embryogenesis by means of Single Cell RNA-Sequencing (scRNA-seq), early mouse embryos were dissected at E5.5 and cultured overnight in the presence or absence of caspase inhibitors. On the following morning, embryos were dissociated with Accutase and subjected to single-cell sorting into 384-well plates. Total RNA isolation was then carried out using a RNA extraction Kit (RNeasy Mini Kit, Qiagen). scRNA-seq was performed using the Smart-seq2 protocol^68^. PolyA selection/enrichment with Ultra II Kit (NEB) was the method adopted for library preparation.

### Data processing, quality control and normalization

We performed transcript quantification in our scRNA-seq data by running Salmon v0.8.2 ^69^ in the quasi-mapping-based mode. First, a transcriptome index was created from the mouse reference (version GRCm38.p4) and ERCC spike-in sequences. Then, the quantification step was carried out with the “quant” function, correcting for the sequence-specific biases (“--seqBias” flag) and the fragment-level GC biases (“--gcBias” flag). Finally, the transcript level abundances were aggregated to gene level counts. On the resulting raw count matrix including 1,495 cells, we apply a quality control to exclude poor quality cells from downstream analyses.

For the quality control we used the following criteria: we identified the cells that have a log_10_ total number of reads equal to or greater than 4, a fraction of mapped reads equal to or greater than 0.8, a number of genes with expression level above 10 reads per million equal to or greater than 3000 and a fraction of reads mapped to endogenous genes equal to or greater than 0.5. This resulted in the selection of 723 cells, which were kept for downstream analyses. Transcripts per million (TPM) normalization (as estimated by Salmon) was used.

### Identification of highly variable genes and dimensionality reduction

To identify highly variable genes (HVG), first we fitted a mean-total variance trend using the R function “trendVar” and then the variance was decomposed into biological and technical components with the R function “decomposeVar”; both functions are included in the package “scran” (version 1.6.9 ^70^).

We considered HVGs those that have a biological component that is significantly greater than zero at a false discovery rate (Benjamini-Hochberg method) of 0.05. Then, we applied further filtering steps by keeping only genes that have an average expression greater to or equal than 10 TPM and are significantly correlated with one another (function “correlatePairs” in “scran” package, FDR<0.05). This yielded 1921 genes, which were used to calculate a distance matrix between cells defined as 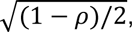, where ρ is the Spearman’s correlation coefficient between cells. A 2D representation of the data was obtained with the UMAP package (version 0.2.0.0 https://cran.r-project.org/web/packages/umap/index.html) using the distance matrix as input.

### Cell clustering and connectivity analysis

To classify cells into different clusters, we ran hierarchical clustering on the distance matrix (see above; “hclust” function in R with ward.D2 aggregation method) followed by the dynamic hybrid cut algorithm (“cutreeDynamic” function in R package “dynamicTreeCut” (https://CRAN.R-project.org/package=dynamicTreeCut) version 1.63.1, with the hybrid method, a minimum cluster size of 35 cells and a “deepSplit” parameter equal to 0), which identified five clusters. Cells from different batches were well mixed across these five clusters (see Extended Data Fig. 1), suggesting that the batch effect was negligible. The identity of the five clusters was established based on the expression of known marker genes of Epiblast, Visceral Endoderm and Extra-Embryonic Ectoderm, which were identified in a previous study^56^. The expression levels of some of the top markers is plotted in Figure 1b.

We performed a robustness analysis on the clustering by exploring in detail how the choices of genes, clustering parameters and algorithms affect the identity and the number of clusters. First, we quantified the cluster robustness by calculating Pearson’s gamma and the Average silhouette width obtained with 100 random subsets of 60% of the highly variable genes and different values of the deepSplit parameter. While the robustness at deepSplit=0 and 1 is similar, for greater values of deepSplit (corresponding to less conservative clustering) the robustness rapidly declines (Extended Data Figure 2a). The clustering with deepSplit = 0 and 1 (the more robust choices) yield very similar results, the only difference being the splitting of the intermediate cluster in two subclusters (Extended Data Figure 2b).

In addition to this, we also used Louvain clustering on the highly variable genes (resolution=0.3, k=20 with 20 principal components), which again produced very similar clusters.

We quantified the connectivity between the clusters (using only CI-treated cells) with PAGA^22^ implemented in the python library scanpy (version 1.4.7)^71^. The analysis revealed that the three epiblast clusters are connected with each other while the two extra embryonic tissues (Visceral Endoderm and Extra Embryonic Ectoderm) are isolated (Extended Data Figure 2c).

### Identification of a single-cell trajectory in the epiblast

We calculated a diffusion map (“DiffusionMap” function in the R package “destiny” version 2.6.2 ^23^ on the distance defined above on the epiblast cells from CI-treated embryos. The pseudotime coordinate was computed with the “DPT” function with the root cell in the winner epiblast cluster (identified by the function “tips” in the “destiny” package). Such pseudotime coordinate can be interpreted as a “losing score” for all the epiblast cells from the CI-treated embryos.

We estimated the losing scores of the epiblast cells from DMSO-treated embryos by projecting such data onto the diffusion map previously calculated (function “dm_predict” in the destiny package). Finally, for each of the projected cells, we assigned the losing score as the average of the losing scores of the 10 closest neighbours in the original diffusion map (detected with the function “projection-dist” in the destiny package).

While for the clustering and the trajectory analysis we used the highly variable genes computed from the whole dataset, we verified that all results concerning the separation between winner and loser epiblast cells (eg, clusters and losing score) remain unaffected if the highly variable genes are calculated using only the epiblast cells.

### Mapping of data from epiblast cells onto published single-cell RNA seq datasets of epiblast from freshly isolated embryos

We compared the transcriptional profile of epiblast from embryos cultured in DMSO and CI with that of epiblast collected from freshly isolated embryos at different stages.

To do this, we considered the dataset published in^26^, which includes epiblast cells from embryos at the stages E5.5 (102 cells), E6.25 (130 cells) and E6.5 (288 cells). A diffusion map and a diffusion pseudotime coordinate were computed with these cells following the same procedure described in the section above (Extended Data Figure 2d-e). Then, we projected epiblast cells from CI and DMSO-treated embryos and we assigned to them a diffusion pseudotime coordinate as described above (Extended Data Figure 2f).

### Differential gene expression analysis along the trajectory

To identify the genes that are differentially expressed along the trajectory, first we kept only genes that have more than 15 TPM in more than 10 cells (this list of genes is provided in Supplementary Table 4); then, we obtained the log-transformed expression levels of these genes (adding 1 as a pseudo-count to avoid infinities) as a function of the losing score and we fitted a generalized additive model to them (R function “gam” from “GAM” package version 1.16.). We used the ANOVA test for parametric effect provided by the gam function to estimate a p-value for each tested gene. This yielded a list of 5,311 differentially expressed genes (FDR < 0.01).

Next, we looked for groups of differentially expressed genes that share similar expression patterns along the trajectory. To this aim, similarly to what we did when clustering cells, we calculated a correlation-based distance matrix between genes, defined as 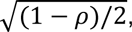, where ρ is the Spearman’s correlation coefficient between genes. Hierarchical clustering was then applied to this matrix (hclust function in R, with ward.D2 method) followed by the dynamic hybrid cut algorithm (dynamicTreeCut package) to define clusters (“cutreeDynamic” function in R with the hybrid method and a minimum cluster size of 100 genes and a deepSplit parameter equal to 0). This resulted in the definition of four clusters, three of genes that decrease along the trajectory (merged together for the GO enrichment and the IPA analysis) and one of increasing genes (Extended Data Fig. 2d). IPA (QIAGEN Inc., https://www.qiagenbio-informatics.com/products/ingenuity-pathway-analysis), was run on all genes differentially expressed (FDR < 0.01) along the trajectory from winner to loser cells (see Fig. 2a-d and Fig. 3a-c), using all the tested genes as a background (see Supplementary Table 4). This software generated networks, canonical pathways and functional analysis. The list of decreasing/increasing genes is provided in Supplementary Tables 1 and 2.

#### Analysis of Mitochondrial DNA heteroplasmy in single-cell RNA seq dataset

We used STAR (version 2.7 ^72^) to align the transcriptome of the epiblast cells from CI-treated embryos (274) to the mouse reference genome (mm10). Only reads that uniquely mapped to the mitochondrial DNA (mtDNA) were considered. From these, we obtained allele counts at each mtDNA position with a Phred Quality Score greater than 33 using the samtools mpileup function. Next, we applied filters to remove cells and mtDNA positions with a low coverage. First, we removed cells with fewer than 2,000 mtDNA positions covered by more than 50 reads. Second, we removed positions having less than 50 reads in more than 50% of cells in each of the three epiblast clusters (winner, intermediate and loser). These two filters resulted in 259 cells and 5,192 mtDNA positions (covered by ∼700 reads per cell on average) being considered for further analyses.

Starting from these cells and positions, we applied an additional filter to keep only positions with a sufficiently high level of heteroplasmy. To this aim, for each position with more than 50 reads in a cell, we estimated the heteroplasmy as:

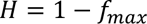

where *f_max_* is the frequency of the most common allele. We kept only positions with *H*>0.01 in at least 10 cells.

Finally, using generalized additive models (see above), we identified the positions whose heteroplasmy *H* changes as a function of the cells’ losing score in a statistically significant way. We found a total of eleven significant positions (FDR < 0.001), six of them in the *mt-Rnr1* gene and five in the *mt-Rnr2* gene. All of these positions have a higher level of heteroplasmy in loser cells (see Fig. 6b-g and Extended Data Fig. 6f-k). The results remain substantially unaltered if the Spearman’s rank correlation test (in alternative to the generalized additive models) is used. For the barplot shown in Fig. 6h and the correlation heatmaps in Fig. 6i and Extended Data Fig. 6l, we took into account only cells that covered with more than 50 reads all the significant positions in the *mt-Rnr1* gene (215 cells, Fig. 6h-6i) or in both the *mt-Rnr1* and *mt-Rnr2* genes (214 cells, Extended Data Fig. 6l).

As a negative control, we repeated the analysis described above using the ERCC spike-ins added to each cell. As expected, none of the positions was statistically significant, which suggested that our procedure is robust against sequence errors introduced during PCR amplification.

We also performed the mtDNA heteroplasmy analysis in cells from the Visceral Endoderm and the Extra-Embryonic Ectoderm in both DMSO and CI conditions: none of these cells have a mtDNA heteroplasmy higher than 0.01 in the 11 significant positions identified within mt-Rnr1 and mt-Rnr2 in loser epiblast cells, and the reference allele is always the most common. This reinforces the hypothesis that such variants are specific to loser epiblast cells and are not resulting from contamination.

To test the reliability of our heteroplasmy estimations, we used the RNA-seq data from two of the mtDNA cell lines (BG and HB, see Figure 7), for which the heteroplasmy was measured also by ARMS-qPCR. To do so, first we downloaded the fasta files of the two mtDNA cell lines from https://www.ncbi.nlm.nih.gov/nuccore/KC663619.1 and https://www.ncbi.nlm.nih.gov/nuccore/KC663620.1, then we identified the mtDNA positions that differ from the BL6 reference genome. Finally, on these different positions, the heteroplasmy *H* was computed as explained above. The values of heteroplasmy we found with our computational analysis were very close to those estimated by ARMS-qPCR (∼17% from RNA-seq data vs ∼21% measured by ARMS-qPCR; and ∼93% from RNA-seq data vs ∼97% by measured by ARMS-qPCR).

### Common features of scRNA-seq and bulk RNA-seq datasets

Differential expression analysis between the co-cultured winner HB(24%) and loser cell line BG(95%) was performed using the package EdgeR version 3.20.9 ^73^.

Batches were specified in the argument of the function model.matrix. We fitted a quasi-likelihood negative binomial generalized log-linear model (with the function glmQLFit) to the genes that were filtered by the function filterByExpr (with default parameter). These genes were used as background for the gene enrichment analysis.

We set a FDR of 0.001 as a threshold for significance. The enrichment analysis for both the scRNA-seq and bulk RNA-seq datasets were performed using the tool g:Profiler ^74^. The list of up-regulated, down-regulated and background genes related to the DE analysis for the bulk RNA-seq dataset are provided in the Supplementary Tables 5, 6 and 7.

### Quantification and Statistical Analysis

Box plots show lower quartile (Q1, 25^th^ percentile), median (Q2, 50^th^ percentile) and upper quartile (Q3, 75^th^ percentile). Box length refers to interquartile range (IQR, Q3-Q1). The upper whisker marks the minimum between the maximum value in the dataset and the IQR times 1.5 from Q3 (Q3+1.5 x IQR), while the lower whisker marks the maximum between the minimum value in the dataset and IQR times 1.5 from Q1 (Q1-1.5 x IQR). Outliers are shown outside the interval defined by box and whiskers as individual points.

Flow cytometry data was analysed with FlowJo Software.

Western blot quantification was performed using Image Studio Lite (LI-COR). Protein expression levels were normalised to loading controls vinculin or α-tubulin.

The quantification of the DDIT3 and OPA1 expression in embryos was done by two distinct methods. DDIT3 expression was quantified by counting the number of epiblast cells with positive staining in the embryos of each group. The expression of OPA1 was quantified on Fiji software as the mean fluorescence across a 10 pixel width line drawn on the basal cytoplasm of each cell with high or low p-rpS6 fluorescence intensity, as specified in^7^. min of 8 cells were quantified per condition (high vs low mTOR activity) in each embryo. Six embryos treated with CI were analysed. Mean grey values of OPA1 fluorescence for each epiblast cell are pooled on the same graph.

Normalisation of data from metabolic flux analysis with Seahorse was performed using Wave Desktop software (Agilent Technologies, UK) and data exported to Prism 8 (GraphPad) for statistical analysis.

The statistical analysis of the results was performed using GraphPad Prism version 8.0.0 for Mac (GraphPad Software, San Diego, California USA). Data was tested for normality using Shapiro-Wilk normality test. Parametric or non-parametric statistical tests were applied accordingly. Details about the test used in each of the experiments are specified in figure legends. Statistical significance was considered with a confidence interval of 0.05%. n.s., non-significant; * p<0.05; ** p<0.01;*** p<0.001.

### Data Availability

Data were analysed with standard programs and packages, as detailed above. Authors can confirm that all relevant data are included in the paper and/ or its supplementary information files. Source data for Figures 2-5,7 and for Extended Data Figures 4-5, 7-8 are provided with the paper. RNA-seq raw as well as processed data are available through ArrayExpress, accession numbers E-MTAB-8640, for scRNA-seq data, and E-MTAB-8692, for bulk RNA-seq data.

### Code Availability

All code that was used in this study is available upon request.

## Notes

### Competing Interest Statement

The authors have declared no competing interest.

